# Chemokine receptor 2-targeted molecular imaging in pulmonary fibrosis

**DOI:** 10.1101/2020.03.04.960179

**Authors:** Steven L. Brody, Sean P. Gunsten, Hannah P. Luehmann, Debbie H. Sultan, Michelle Hoelscher, Gyu Seong Heo, Jiehong Pan, Jeffrey R. Koenitzer, Ethan C. Lee, Tao Huang, Cedric Mpoy, Shuchi Guo, Richard Laforest, Amber Salter, Tonya D. Russell, Adrian Shifren, Christophe Combadiere, Kory J. Lavine, Daniel Kreisel, Benjamin D. Humphreys, Buck E. Rogers, David S. Gierada, Derek E. Byers, Robert J. Gropler, Delphine L. Chen, Jeffrey J. Atkinson, Yongjian Liu

**Affiliations:** Department of Medicine, Washington University School of Medicine, Saint Louis, MO USA; Department of Radiology, Washington University School of Medicine, Saint Louis, MO USA; Department of Radiation Oncology, Washington University School of Medicine, Saint Louis, MO USA; Division of Biostatistics, Washington University School of Medicine, Saint Louis, MO USA; Department of Developmental Biology, Washington University School of Medicine, Saint Louis, MO USA; Department of Surgery, Washington University School of Medicine, Saint Louis, MO USA; Department of Immunology and Pathology, Washington University School of Medicine, Saint Louis, MO USA; Sorbonne Université, Inserm, CNRS, Centre d’Immunologie et des Maladies Infectieuses, Cimi-Paris, F-75013, Paris, FR

## Abstract

Idiopathic pulmonary fibrosis (IPF) is a progressive, inflammatory lung disease that is monitored clinically by measures of lung function, without effective molecular markers of disease activity or therapeutic efficacy. Lung immune cells active in the pro-fibrotic process include inflammatory monocyte and interstitial macrophages that express the C-C motif chemokine receptor 2 (CCR2). CCR2^+^ monocyte lung influx is essential for disease phenotypes in models of fibrosis and identified in lungs from subjects with IPF. Here, we show that our peptide-based radiotracer ^64^Cu-DOTA-ECL1i identifies CCR2^+^ inflammatory monocytes and interstitial macrophages in multiple preclinical mouse models of lung fibrosis, using positron emission tomography (PET) imaging. Mice with bleomycin-induced fibrosis treated with blocking antibodies to interleukin-1β, a mediator of fibrosis associated with CCR2^+^ cell inflammation, or with pirfenidone, an approved anti-fibrotic agent, demonstrated decreased CCR2-dependent interstitial macrophage accumulation and reduced ^64^Cu-DOTA-ECL1i PET uptake, compared to controls. Lung tissues from patients with fibrotic lung disease demonstrated abundant CCR2^+^ cells surrounding regions of fibrosis, and an ex vivo tissue-binding assay showed correlation between radiotracer localization and CCR2^+^ cells. In a phase 0/1 clinical study of ^64^Cu-DOTA-ECL1i PET, healthy volunteers showed little lung uptake, while subjects with pulmonary fibrosis exhibited increased uptake, notably in zones of subpleural fibrosis, reflecting the distribution of CCR2^+^ cells in the profibrotic niche. These findings support a pathologic role of inflammatory lung monocytes/macrophages in fibrotic lung disease and the translational use of ^64^Cu-DOTA-ECL1i PET to track CCR2-specific inflammation for image-guided therapy.

**One Sentence Summary:** PET imaging of CCR2^+^ cells in lung fibrosis identifies a therapeutic response in mouse models and displays a perifibrotic signal in subjects with IPF.

## Introduction

Idiopathic pulmonary fibrosis (IPF) is a devastating lung disease characterized by interstitial macrophage accumulation, fibroblast proliferation, and matrix deposition of uncertain pathogenesis (*1*). Patients have a clinical course that ranges from slow progression to rapid deterioration and an overall five-year survival of 20 to 40% (*2*). Anti-fibrotic therapies are limited; pirfenidone and nintedanib are currently the only approved medications for IPF (*3, 4*). These drugs slow disease progression but have a limited positive impact on survival (*3, 4*). However, there is currently no way to predict the individual patient’s response to a specific therapy, nor are there established markers to monitor the molecular or cellular response to a treatment. Particular to IPF, bronchoalveolar lavage, and especially lung biopsy, are often avoided due to clinical status and a tendency to worsen disease (*5, 6*). Thus, the development of a non-invasive, molecular assessment may address these challenges to improve patient care and the therapeutic pipeline.

A major gap in IPF care remains a paucity of tools to define patient-specific molecular phenotypes, including lung or serum markers to follow the course of disease. Substantial efforts to develop non-invasive testing, such as genetic signatures in peripheral leukocytes, have been imprecise, though elevated levels of circulating CD14^+^ monocytes are recently suggested to predict mortality (*7*). Clinically, features of fibrosis present on high resolution chest computed tomography (CT) are used for diagnosis and can predict mortality (*5, 6*). The primary biomarkers used to follow disease are pulmonary function test parameters: forced vital capacity (FVC), diffusion capacity for carbon monoxide, and the distance walked in 6 minutes (*5, 8*). While these tests validate changes in pulmonary physiology, they are indirect measures of disease outcome (*3, 6, 8*). Most physiologic measures do not reverse with effective therapeutics and instead primarily reflect irreversible remodeling rather than a measure of profibrotic activity.

CCR2^+^ (C-C motif chemokine receptor 2) cells have a context-dependent activity in the fibrotic lung and represent a rational marker of inflammation in fibrotic disease. In vivo studies show that elevated levels of lung tissue CCL2 (C-C motif chemokine ligand 2) contribute to lung fibrosis by directing Ly6C^high^, CCR2^+^ inflammatory (classical) monocyte egress from the bone marrow to the lung (*9, 10*). *Ccr2*-deficient mice have markedly attenuated development of lung fibrosis induced by bleomycin, radiation, and other pro-fibrotic irritants (*11–13*). In lung fibrosis models, lineage tracing and single cell transcriptome analysis show that CCR2^+^ monocytes accumulate in the lung and differentiate into cells referred to as inflammatory monocyte-derived macrophages, interstitial macrophages, or tissue macrophages (*10, 14, 15*). In humans, CCR2-dependent bone marrow-derived monocytes and interstitial macrophages are increased in samples taken from the lungs of patients with pulmonary fibrosis (*15–17*). CCR2^+^ monocytes and macrophages produce factors that induce fibroblast recruitment and collagen production including TGFβ, implicating the interstitial macrophages in the pro-fibrotic process (*14–21*). However, the temporal infiltration and differentiation of CCR2^+^ monocytes relative to degree of inflammation or fibrosis is not well known in experimental models nor is their distribution in human fibrotic lungs well described. Moreover, scant detail exists regarding the therapeutic effects of existing anti-fibrotic drugs on CCR2^+^ and CCR2-derived populations (*22, 23*). A detailed map of CCR2^+^ cells in the fibrotic lung would be essential for evaluating the role of CCR2 in emerging molecular-based therapies.

Recently, our group reported on the development of a peptide-based radiotracer, ^64^Cu-DOTA-ECL1i, to quantify the CCR2-specific inflammatory cell burden using positron emission tomography (PET) in pre-clinical models. The sensitivity and specificity of ^64^Cu-DOTA-ECL1i for imaging CCR2^+^ cell trafficking was demonstrated in experimental lung injury induced by endotoxin or reperfusion, atherosclerosis, and myocardial injury in mice (*24–27*). Application of this radiotracer for clinical use in patients with pulmonary fibrosis is attractive based on the requirement for CCR2^+^ immune cells in experimental models and their presence in human diseased lungs. Also compelling are the clinical challenges of managing patients with IPF in the face of a paucity of actionable molecular markers of disease activity. Accordingly, we used mouse models to study the late stages of lung fibrosis induced by bleomycin and ionizing radiation to establish that increased ^64^Cu-DOTA-ECL1i PET uptake in the lung correlates with CCR2^+^ cell infiltration associated with fibrosis. We then demonstrated that therapeutic modulation of fibrosis by using anti-IL-1β to block activity in interstitial macrophages or treatment with the anti-fibrotic drug pirfenidone, decreased the PET uptake signal to provide a unique clinically applicable measure. We advanced the radiotracer to first-in-human use in patients with pulmonary fibrosis to reveal increased uptake in regions of subpleural fibrosis in the lung. The findings suggest the clinical relevance of CCR2 as a molecular target for pulmonary fibrosis and support the translational use of ^64^Cu-DOTA-ECL1i to monitor CCR2-specific inflammatory cell activity. Future applications may facilitate the development of targeted therapies that alter CCR2-mediated profibrotic remodeling.

## Results

### CCR2^+^ cells associate with perifibrotic regions in lungs of mice with bleomycin-induced fibrosis

CCR2^+^ cells are tightly linked with the development of lung fibrosis as indicated by studies in *Ccr2^−/−^* mice, however the temporal and spatial nature of the CCR2^+^ cells and their progeny relative to fibrosis is not well described. To track CCR2^+^ cells types during development of bleomycin-induced fibrosis, we used a transgenic *Ccr2^gfp/+^* knockin/knockout reporter mice harboring the enhanced green fluorescent protein (EGFP) sequence in the *Ccr2* gene. *Ccr2^gfp/+^* and *Ccr2* null (*Ccr2^gfp/gfp^*) mice were intranasally administered bleomycin as an established but imperfect model of IPF (*28–30*). In this model, inflammation rises on days 3 through 10 post-bleomycin, followed by the development of fibrosis over 14 to 28 days. Serial tissue sections were scored for EGFP-expressing cells and fibrosis in merged images (**Fig. 1A-C; Fig. S1**). Compared to control mice, the numbers of CCR2-EGFP^+^ cells in lung sections were significantly increased at 14 days after bleomycin delivery, then diminished at day 28 (**Fig. 1B**). CCR2-EGFP^+^ cell incidence mapped with fibrosis quantified by the modified Ashcroft score (*31*) (**Fig. 1C**). The abundance of CCR2-EGFP^+^ cells was relatively low in regions of both normal appearing lung and dense fibrosis. Instead, these cells were abundant in regions that surrounded the remodeled parenchyma and those areas near fibrosis, supportive of a role for CCR2^+^ cells in the fibrotic niche. At days 14 and 28 there was significantly greater accumulation of CCR2-EGFP^+^ cells at the highest Ashcroft scores (**Fig. 1C**).

**Figure 1.**
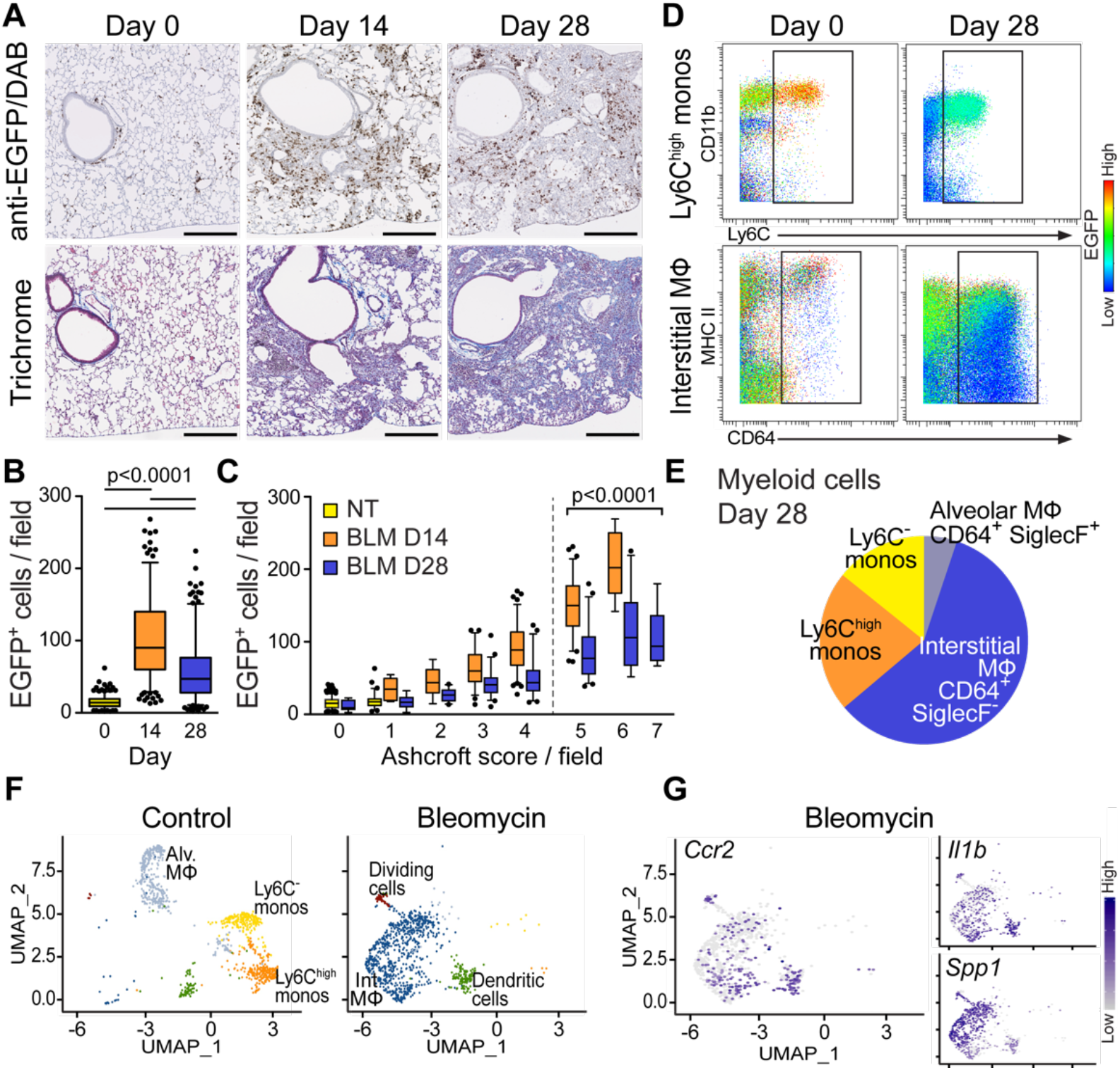
CCR2^+^ cells localize to perifibrotic regions in bleomycin-induced lung fibrosis. C57BL/6 *Ccr2^gfp/+^* mice were administered intranasal bleomycin and lungs assayed at the indicated day. (**A**) Representative images of CCR2-EGFP^+^ cells, identified using an anti-EGFP antibody (brown, top), with serial tissue sections stained using trichrome (bottom). (**B**) Quantitation of CCR2-EGFP^+^ cells in lung sections (0.14 mm^2^ /field; approximately 250 fields/lung, n=3-5 mice/time point). (**C**) CCR2-EGFP^+^ cells in lung sections at each modified Ashcroft fibrosis score. Each cell count and fibrosis score are from the same field in serial sections. Significantly more CCR2-EGFP^+^ cells were in the regions with Ashcroft scores 5-7 than 0-4 at 14 and 28 days. (**D**) Representative pseudo-colored density plots from analysis of mass cytometry showing a shift in the percentage of CCR2-EGFP^+^ inflammatory monocytes (Ly6C^high^ monos) and interstitial macrophages (MФ) in single cell preparations of whole mouse lungs, days 0 and 28 post-bleomycin. (**E**) Relative proportions of CCR2-EGFP^+^ myeloid cell populations at day 28 post-bleomycin identified by mass cytometry (n=5 mice/day). Cells in D and E were identified using the gating strategy in Supplementary Methods and Fig. S2. (**F**) Representative differences in transcription of myeloid single-cell populations isolated from lungs of wild-type mice (day 20). Single-cell RNA sequencing of cells was visualized using a uniform manifold approximation and projection (UMAP) plot. Cell types clustered by identity. (**G**) Cells expressing *Ccr2* proinflammatory and profibrotic genes in interstitial macrophages post bleomycin. Significance in A and B was determined by Kruskal-Wallis with Dunn’s Multiple Comparison test. In A, Bars=250 μm.

Mass cytometry was used to characterize types of CCR2^+^ cells found in bleomycin-induced injury. A cocktail of 35 cell marker antibodies (**Table S1**) allowed for accurate classification of immune cell populations on a single cell basis, as identified previously in the bleomycin model (**Fig. 1D**; **Fig. S2A**) (*32, 33*). At baseline, low numbers of CCR2-EGFP^+^ cells (**Fig 1A,B**) with characteristic markers of monocytes, dendritic cells and rare lymphocytes were present. Following bleomycin delivery, the percentage of GFP^+^ myeloid cells markedly increased, in concert with CCR2^+^ inflammatory monocytes. At day 28 post bleomycin, interstitial macrophage, which are derived from CCR2^+^, Ly6C^high^ inflammatory monocytes (*15, 32, 34*), were the most abundant EGFP^+^ myeloid cells type (**Fig. 1D, E, Fig. S2B**). As expected, *Ccr2^gfp/gfp^* mice lacked inflammatory monocytes and interstitial macrophages at day 0 and 28 (**Fig. S2B**). To confirm the observed shift in monocyte-to-macrophage cell types and further characterize the CCR2-expressing cells, we performed single cell RNA sequencing of the mouse lung at day 20 post-bleomycin, providing high precision of immune cell phenotypes, while agnostic to current surface markers (*14, 15*). In control lungs, *Ccr2* expression primarily clustered in a subset of monocyte-lineage cells that co-expressed inflammatory monocyte and dendritic cell genes (**Fig. 1F, G**). After bleomycin, *Ccr2*^+^ interstitial macrophages were enriched in genes previously associated with pro-fibrosis pathways in mouse models and human lung tissues, notably *Il-1b*, and *Spp1*, *Apoe, Mmp14*, and others (**Fig. 1G, S3**) (*14–16, 19, 35*). These results describe a temporal and spatial relationship of CCR2-expressing cells with the development of a fibrotic niche and are consistent with the reported role of CCR2^+^ cells in genetic models.

### ^64^Cu-DOTA-ECL1i PET lung uptake is increased during bleomycin fibrosis

We have previously developed ^64^Cu-DOTA-ECL1i to image CCR2^+^ cells by PET and reported the accuracy of this radiotracer uptake in preclinical models (*24, 25, 27*). As a next step, we examined ^64^Cu-DOTA-ECL1i and PET/CT application in lung fibrosis using the bleomycin-injury mouse model. The uptake of radiotracer in the lungs of mice given bleomycin, compared to controls, showed minimal increase at day 2, with significant increase at day 14 (**Fig. 2A-D**). Uptake diminished at day 28, though remained significantly above control levels. The changes in PET signal were comparable to the relative abundance of CCR2^+^ cells detected by immunostaining at days 0, 14 and 28 (**Fig. 1B, Fig. 2D**). Molecular specificity of ^64^Cu-DOTA-ECL1i for detection of CCR2^+^ cell burden was demonstrated by the significantly decreased uptake in *Ccr2^gfp/gfp^* knockout mice compared to wild-type mice at day 14 (**Fig. 2B**). Radiotracer specificity was confirmed by decreased uptake in mice injected with non-radioactive ECL1i as competitive receptor blockade (**Fig. 2C**). To approximate the relationship of the ^64^Cu-DOTA-ECL1i PET signal and regions of fibrotic remodeling, fixed lungs were sectioned through the coronal plane after scanning. Trichrome-stained lung sections aligned with the PET images showed prominent ^64^Cu-DOTA-ECL1i uptake in regions of lung inflammation and fibrosis (**Fig. 2E**). The relationship of radiotracer uptake and fibrosis was consistent with an enrichment of CCR2-EGFP^+^ cells in perifibrotic regions of the lung. These data suggest that ^64^Cu-DOTA-ECL1i PET uptake mirrors regional differences in CCR2^+^ cell recruitment during inflammation and the progression of fibrosis.

**Fig. 2.**
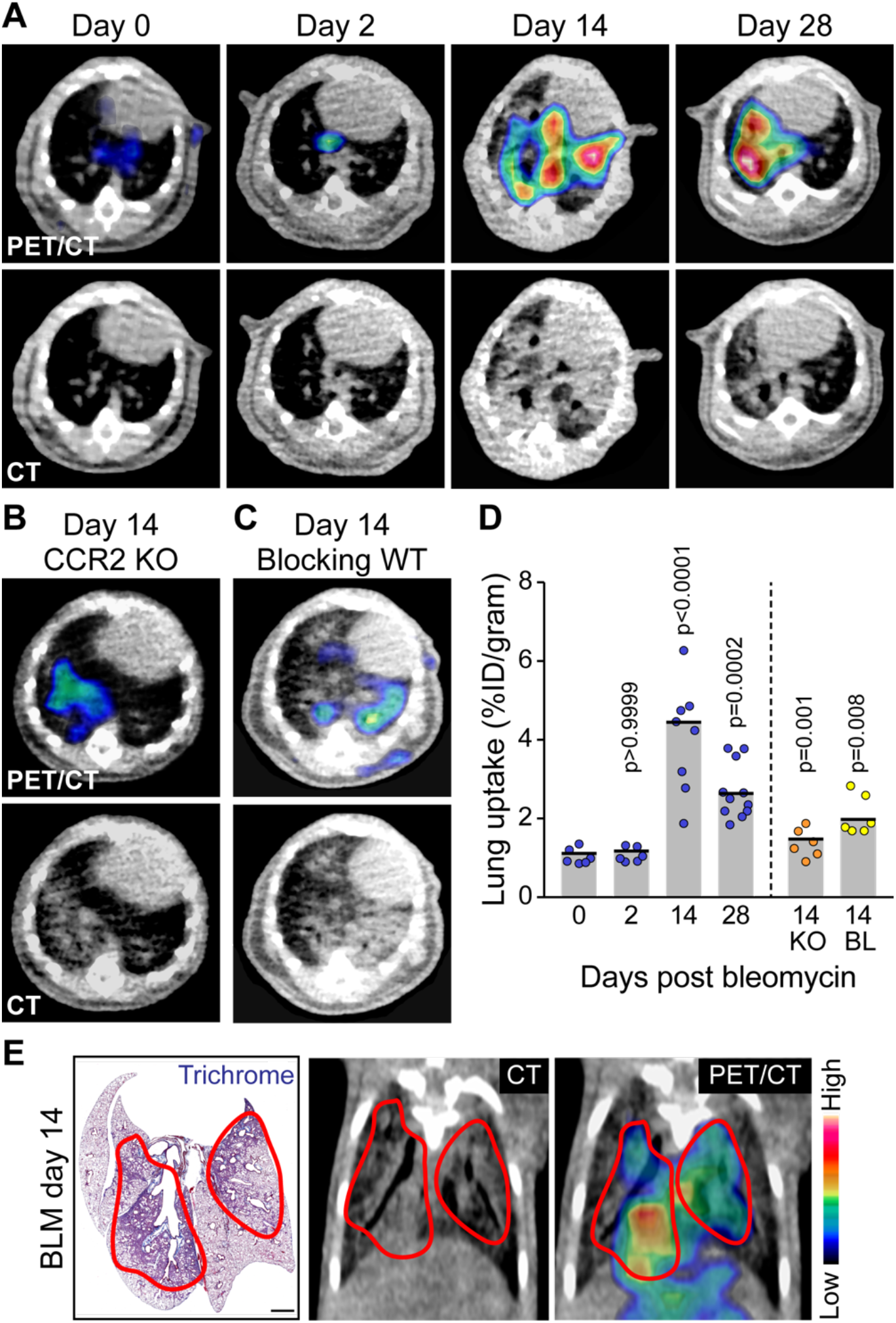
Detection of ^64^Cu-DOTA-ECL1i uptake by PET/CT in bleomycin-induced lung fibrosis in mice. **A-D**. Mice given intranasal bleomycin were injected with ^64^Cu-DOTA-ECL1i, prior to dynamic PET/CT imaging on the indicated day. Representative transverse images are shown of (**A**) Wild-type (WT), (**B**) *Ccr2* knockout (KO, CCR^gfp/gfp^) at day 14, and (**C**) WT in competitive receptor studies (Blocking) at day 14. (**D**) Lung uptake from A-C. Shown are the medians of n=6-10 mice/day of mixed sexes compared to non-treated mice by Kruskal-Wallis with Dunn’s Multiple Comparison test. Day 14 knockout (KO) mice uptake and blocking (BL) was compared to day 14 control conditions using the Mann-Whitney U test. (**E**) Example of regional cellularity and fibrosis (red borders) in a trichrome-stained tissue section compared to CT and PET/CT-fused coronal sections at day 14 post bleomycin. In E, Bar=1000 μm.

### ^64^Cu-DOTA-ECL1i PET uptake is increased in the lungs of mice with radiation-induced fibrosis

Although the responses of monocytes and macrophages in bleomycin-induced fibrosis have been extensively characterized, the subacute, high-inflammatory condition could bias immune cell populations. As an alternative, we examined ^64^Cu-DOTA-ECL1i PET lung activity in a mouse model of radiation-induced injury and fibrosis. Radiation injury is a chronic, indolent process that is accompanied by inflammatory monocytes and interstitial macrophages, increased IL-1β, and mitigated by genetic deletion of *Ccr2* or *Il-1b* in mice (*13, 36*). We ascertained ^64^Cu-DOTA-ECL1i PET uptake in mice after receiving focused, right-lung irradiation with the non-irradiation left lung assigned as a control (**Fig. S4A**). At 14 and 26 weeks after irradiation, radiotracer uptake was significantly increased in the right lung, while little was detected in the left (**Fig. S4B-D**). Similar to the bleomycin fibrosis model, the ^64^Cu-DOTA-ECL1i PET signal was present in regions of fibrotic remodeling, as indicated by alignment of whole lung preparations stained with trichrome with the PET images (**Fig. S4B, C**). At 14 and 26 weeks, there were significantly more CCR2^+^ cells in the irradiated right compared to control left lung (**Fig. S4E**). The findings are consistent with the known *Ccr2*-dependent responses in radiation-induced fibrosis.

### IL-1β blockade of bleomycin-induced fibrosis decreases ^64^Cu-DOTA-ECL1i PET uptake

As a viable clinical marker, ^64^Cu-DOTA-ECL1i PET should also detect a therapeutic modulation of CCR2^+^ inflammatory and pro-fibrotic processes. IL-1β is a proinflammatory cytokine produced by the inflammasome and active in lung fibrosis (*19, 37*). IL-1β is induced by lung delivery of profibrotic agents, can induce fibrosis upon delivery to the mouse airway, and activates CCL2-mediated chemotaxis, leading to increased lung interstitial macrophages (*19, 37–39*). Genetic deficiency of the IL-1 axis reduces inflammation following bleomycin injury (*38, 39*). As expected, interstitial macrophages expressed IL-1β in the bleomycin model (**Fig. S3**). We sought to determine if the bleomycin pro-fibrotic inflammatory process could be interrupted during late inflammation and this response imaged using ^64^Cu-DOTA-ECL1i PET. Mice receiving bleomycin were treated with IgG isotype control or monoclonal antibody against IL-1β from days 10 through 28, a treatment schedule advised for testing antifibrotic therapies in preclinical models (*40*) (**Fig. 3A**). Compared to controls, lung fibrosis was decreased by anti-IL-1β antibody treatment at day 28 (**Fig. 3B,C**). Lung expression of *Ccr2*^+^ cells was also reduced, as detected by mRNA in situ hybridization (**Fig. 3D,E**). In parallel, ^64^Cu-DOTA-ECL1i PET uptake in the lungs of IL-1β antibody treated mice was significantly reduced (**Fig. 3F, G**).

**Fig. 3.**
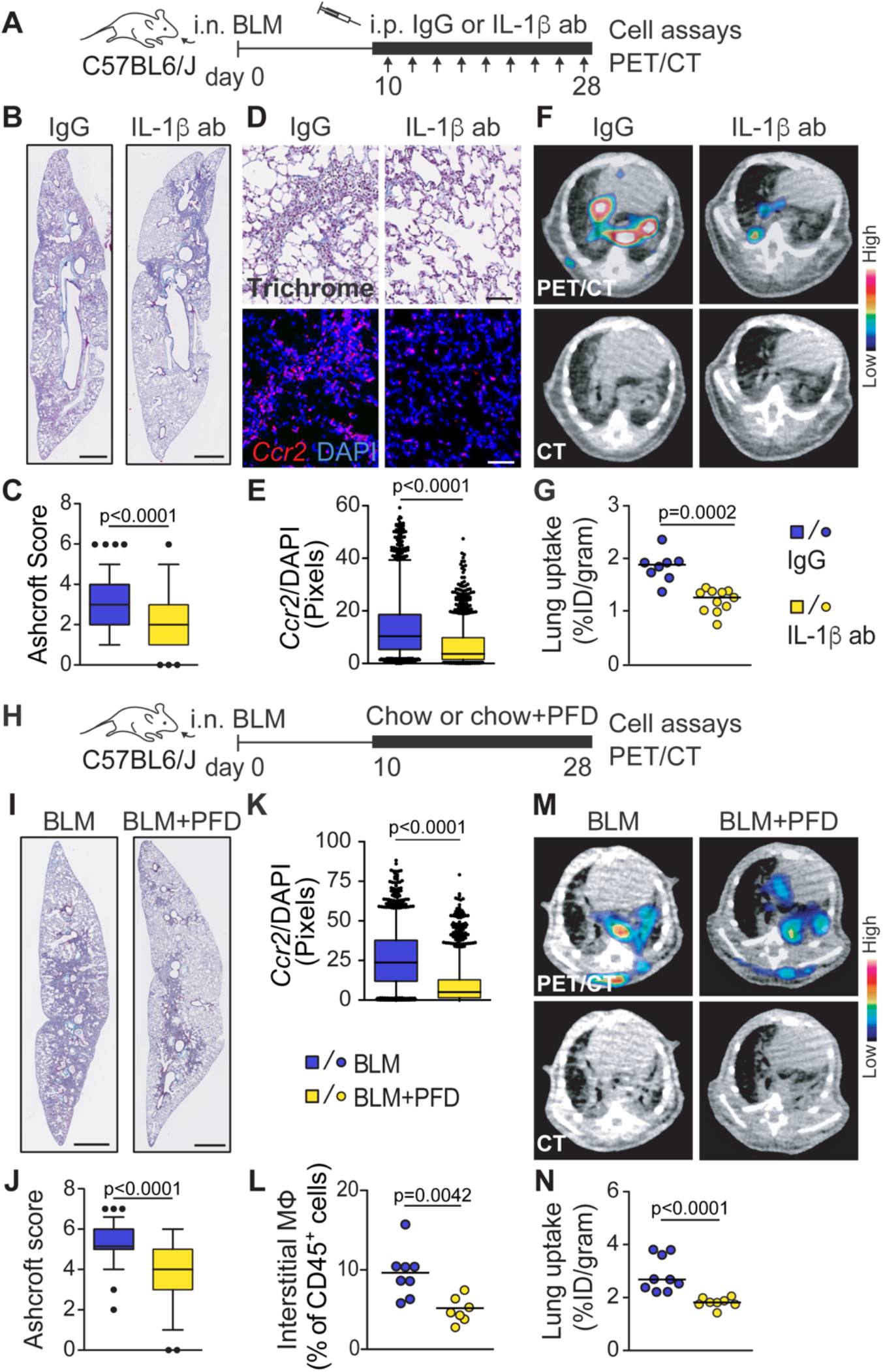
Effect of IL-1β blockade and pirfenidone treatment on ^64^Cu-DOTA-ECL1i PET uptake in bleomycin-induced fibrosis. (**A**) Treatment scheme of wild-type mice administered intranasal bleomycin (i.n. BLM) treated with intraperitoneal (i.p.) IgG or anti-IL-1β antibody three times weekly on days 10-28. **(B**) Representative trichrome-stained lung sections at day 28 (n=3 IgG, 7 IL-1β mice/condition). (**C**) Ashcroft scores from lung sections (0.14 mm^2^ /field; n=360-500 fields/mouse lung; (n=3/group IgG, n=4/group IL-1β). (**D**) Representative images of serial lung tissue sections stained with trichrome (top) and *Ccr2* in situ hybridization (bottom) of indicated condition at day 28. (**E**) Quantitation of *Ccr2* in situ hybridization at day 28 (1400-1900 fields/mouse lung; n=3-4 mice/condition. (**F**) Representative ^64^Cu-DOTA-ECL1i PET/CT uptake in bleomycin-induced lung fibrosis in mice at day 28 treated as indicated. (**G**) PET uptake in lungs after indicated treatment (n=8-9 mice/condition). (**H**) Treatment scheme of mice administered bleomycin, treated with chow only (BLM) or chow containing pirfenidone (BLM+PFD) on days 10-28. (**I**) Representative trichrome-stained lung sections at day 28. (**J**) Ashcroft scores from lung sections (0.14 mm^2^ /field; n=33-36 fields/lung; n=4 mice/condition). (**K**) Quantitation of *Ccr2* in situ hybridization at day 28 (1600-1900, 0.14 mm^2^ fields/mouse lung; n=4 mice/condition). (**L**) Percent of lung interstitial macrophages (MФ, SiglecF^−^/CD64^+^) in single cell preparations of mouse lungs using mass cytometry (n=7-8 mice/condition). (**M**) Representative PET/CT images at day 28 treated as indicated. (**N**) PET uptake in lungs treated as indicated (n=8-9 mice/condition). Significance determined by Mann-Whitney U test for all data. Bars in B, I=1000 μm, D=100 μm.

### Pirfenidone treatment of bleomycin-induced fibrosis decreases ^64^Cu-DOTA-ECL1i PET uptake

While anti-IL-1β antibodies are not a standard IPF treatment, pirfenidone is one of two drugs approved for patients (*3*). Pirfenidone treatment is reported to reduce fibrosis in bleomycin-, radiation-, and graft-versus-host-induced lung fibrosis models (*22, 23, 41, 42*). Thus, we examined pirfenidone modulation of lung ^64^Cu-DOTA-ECL1i PET uptake. Mice were given pirfenidone in chow (*43*) on days 10 through 28 following bleomycin administration (**Fig. 3H**). Compared to mice fed chow only, pirfenidone treatment decreased lung fibrosis (**Fig. 3I, J**), the accumulation of *Ccr2^+^*cells (**Fig. 3K**), and interstitial macrophages (**Fig. 3L**). The treatment effect on CCR2^+^ cell signal was detected as a decrease in ^64^Cu-DOTA-ECL1i PET uptake (**Fig. 3M, N**). These observations indicate that ^64^Cu-DOTA-ECL1i PET can detect changes in CCR2^+^ cells associated with fibrosis and suggest the potential of the PET tracer to monitor therapeutic response in IPF patients or to pre-screen patients for a specific drug treatment.

### CCR2^+^ cells are increased in human fibrotic lung disease

Elevated levels of inflammatory monocytes and tissue/interstitial macrophages are also found in lung tissues of patients with IPF and described by transcriptional profiles (*14, 15, 21, 44, 45*). Characterizing the regional distribution of CCR2^+^ monocytes and macrophages in affected human lung is essential for the clinical application of ^64^Cu-DOTA-ECL1i PET. Certainly, assessing regional differences of CCR2^+^ cellular activity in patients with newly diagnosed or progressive IPF by histology is not feasible. We therefore examined CCR2 expression in lungs removed from patients with end-stage pulmonary fibrosis at lung transplantation (n=11) (**Table S2**) and compared these regions to pre-transplant chest CT images performed for standard clinical evaluation of some subjects. Samples were obtained from regions that included the pleural surface, or within 2 to 5 cm of the pleural edge, using our research protocol for tissue procurement (*24*), which allows mapping and alignment of biopsied regions to chest CT (e.g., samples from the upper lobe, superior segment of the lower lobe). CCR2^+^ cells in lung tissues were identified by immunofluorescent staining of fibrotic tissues and compared to non-fibrotic lungs that were donated but unsuitable for transplantation (control lungs, n=4). Serial sections processed by trichrome staining were used to determine the location of CCR2^+^ cells relative to fibrotic remodeling. CCR2^+^ cells were most abundant in lung tissue sections from subjects with fibrosis compared to no fibrosis (**Fig. 4A, B, C, D**). As in the bleomycin mouse models, CCR2^+^ cells were elevated in cellular regions adjacent to highly fibrotic regions (**Fig. 4A, Fig. S5**), suggesting that regions of high CCR2^+^ cell expression precede the development of fibrotic scar. Mass cytometry was also used to profile CCR2^+^ cell phenotypes in lung biopsies from regions adjacent to the histology sections. In non-fibrotic control lung tissues, CCR2^+^ cells in non-fibrotic lung tissue were primarily CD14^+^, CD16^−^ inflammatory monocytes, as could be predicted in lungs from donors on recent mechanical ventilation (**Fig. S6**). By comparison, there were fewer inflammatory monocytes in the fibrotic lung tissue relative to significant increase in the percent of interstitial macrophages (**Fig. 4E, Fig. S6**).

**Fig. 4.**
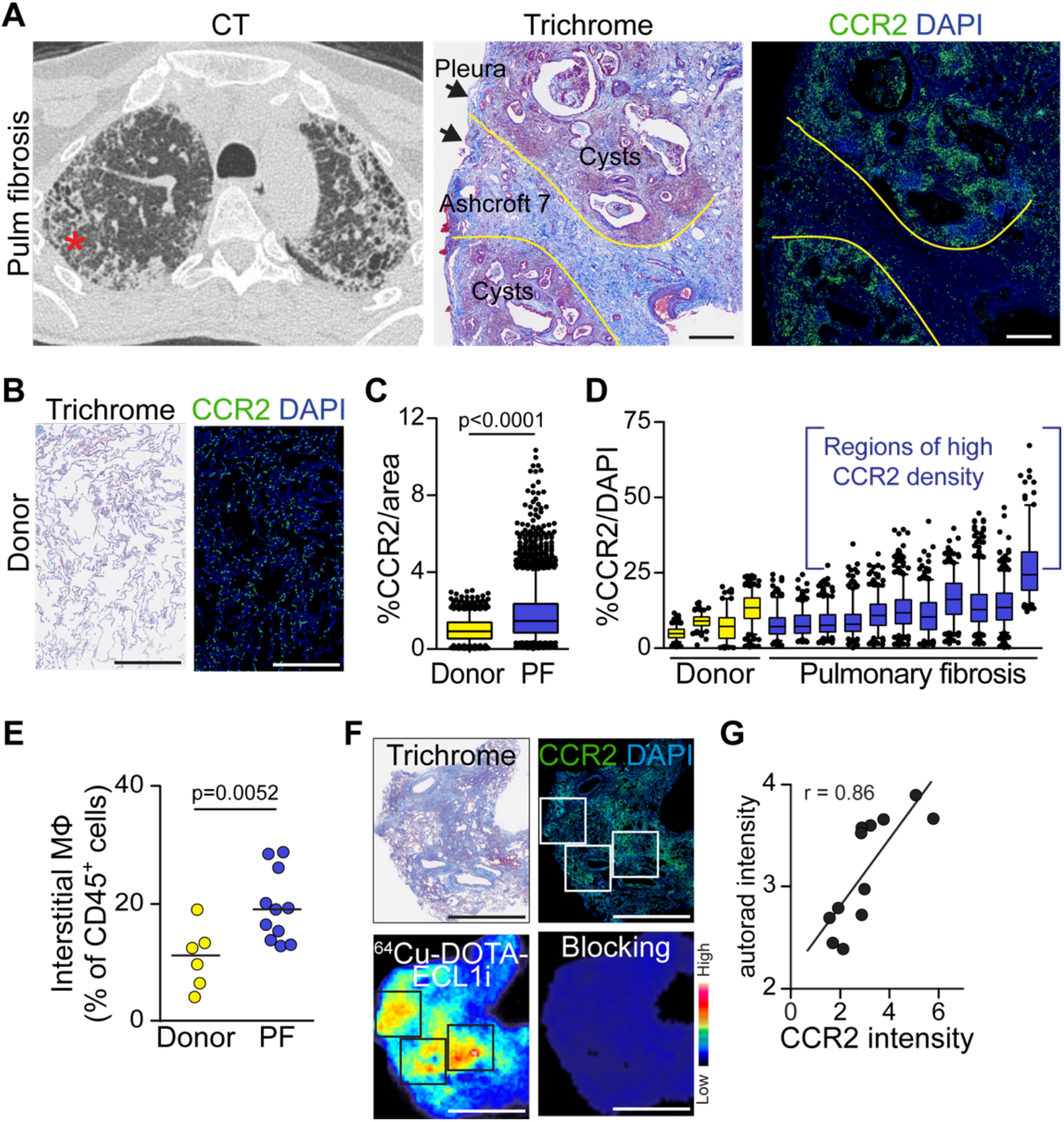
Detection of CCR2^+^ cells in lung tissue explanted from patients with pulmonary fibrosis. Explanted lungs from patients with end-stage pulmonary fibrosis (PF) or non-fibrotic lungs donated for research (Donor). (**A**) Pre-transplant chest CT from a patient with IPF showing the region of explanted lung sampled (*) for serial tissue sections stained by trichrome and CCR2 antibody (green). Arrows indicate pleural surface. Yellow lines demarcate regions of high CCR2^+^ cells. **(B**) Representative serial tissues of donor lung stained as in A. (**C**) Total CCR2^+^ cells in samples (n=4 donors, n=11 PF). (**D**) CCR2^+^ cells from donor and PF lungs. Box plots represent a sample from each subject. Brackets mark fields with CCR2^+^ cell density above the 95th percentile of fields of donors. (**E**) Percent of interstitial macrophages (MФ) in lung tissues (median, n=6 donors, n=11 PF). (**F**) Representative comparison of trichrome, CCR2 staining, and ^64^Cu-DOTA-ECL1i autoradiography in serial sections. Blocking studies confirm binding specificity (n=9). (**G**) Spearman correlation of CCR2 and corresponding ^64^Cu-DOTA-ECL1i pixel intensity in fields of high and low CCR2 immunofluorescence (boxes) from F. In C and E, significance determined by the Mann-Whitney U test. In A,B,E,F DAPI stained nuclei are blue. Bars in A,B=500 μm; F=5 mm.

### ^64^Cu-DOTA-ECL1i uptake is increased in ex vivo lung tissues of subjects with fibrosis

Toward translation to human disease, we next determined if ^64^Cu-DOTA-ECL1i recognized CCR2 in human fibrotic lung tissue (**Table S2**). Lung tissues immunostained for CCR2 were compared to serial sections assayed for ^64^Cu-DOTA-ECL1i by autoradiography (**Fig. 4F, Fig. S7**). Radiotracer specificity was shown by loss of activity in tissues treated with excess non-radioactive ECL1i. Photomicrographs of tissue sections analyzed for CCR2 by immunofluoresent staining and autoradiography were overlaid. Zones of high and low activity of CCR2 staining were compared to ^64^Cu-DOTA-ECL1i binding (**Fig. 4G, Fig. S7**). Although the signal from ^64^Cu autoradiography is lower resolution compared to the high resolution of immunofluorescent microscopy, regional differences in signal were highly concordant. Collectively, the data indicate that CCR2^+^ cells regionally associate with areas of fibrosis in human lung and can be detected ex vivo by ^64^Cu-DOTA-ECL1i.

### ^64^Cu-DOTA-ECL1i PET dosimetry in healthy volunteers

Lung CCR2^+^ cell monitoring may be a valuable clinical tool for the management of patients with pulmonary fibrosis, particularly given the small number of targetable molecular lung markers available. Thus, we obtained approval for use of ^64^Cu-DOTA-ECL1i in humans. The primary aim of this Phase 0/I study was to establish the safety of ^64^Cu-DOTA-ECL1i, to obtain initial estimates of the radiation dosimetry, and to assess lung retention for this marker. Six healthy adult volunteers (3 males, 3 females) with normal lung function were examined (**Table S3**, see human protocol in **Supplementary Methods**). Following intravenous injection of 185 to 370 MBq of ^64^Cu-DOTA-ECL1i, subjects underwent whole body PET/CT scans acquired over about 60 minutes each. Serial PET images were acquired at approximately 30-90 minutes; 2-3 hours; 18-24 hours; and 40-44 hours post injection. Each PET scan was accompanied by a low-dose CT scan for PET attenuation correction. Analysis of images for dosimetry demonstrated minimal lung uptake and predominantly renal clearance (**Fig. 5A, Table S4**). Clearance of ^64^Cu-DOTA-ECL1i from blood was very rapid (**Fig. 5B**). Renal clearance was estimated using urine and bladder activity as a surrogate (**Fig. 5C**). Dosimetry showed the urinary bladder wall as the dose-limiting organ, but overall reasonable dosimetry (**Table S4**). No observable clinically adverse effects of ^64^Cu-DOTA-ECL1i were identified. Low lung uptake and acceptable levels of radiation exposure led us to next evaluate uptake of ^64^Cu-DOTA-ECL1i in the lungs of subjects with pulmonary fibrosis.

**Fig. 5.**
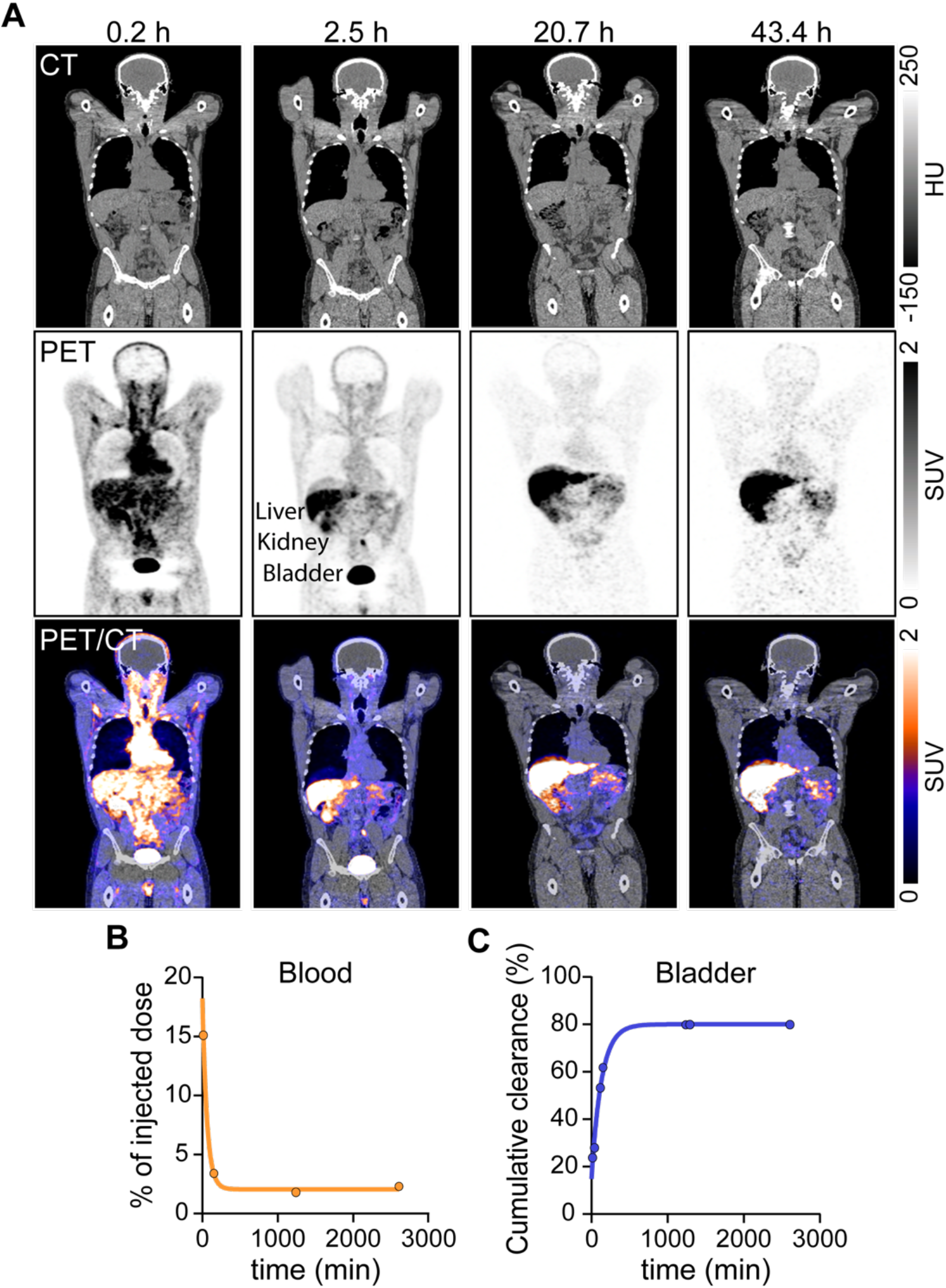
Dosimetry testing of ^64^Cu-DOTA-ECL1i PET/CT imaging in healthy volunteers. (**A**) Representative coronal images of CT, maximum intensity projection PET, and PET/CT from a healthy volunteer obtained at the indicated time following a single intravenous injection with ^64^Cu-DOTA-ECL1i (n=6). PET images show uptake in liver, kidney, and bladder. Representative time-activity curves demonstrate (**B**) rapid blood clearance and (**C**) estimated renal clearance based on changes in bladder activity.

### ^64^Cu-DOTA-ECL1i PET activity is increased in lungs of patients with IPF

Four subjects with IPF underwent ^64^Cu-DOTA-ECL1i PET/CT imaging (**Fig. 6, Table S3**). The pulmonary function testing (FVC) and chest CT features integrated into a fibrotic lung disease score (*46*) that varied among the subjects evaluated (**Table 1**). Subjects with IPF, and an additional healthy volunteer, underwent dynamic PET imaging of the chest for 60 minutes, beginning at the time of intravenous injection of 296 to 370 MBq of ^64^Cu-DOTA-ECL1i. The scan was followed by whole-body images obtained 80-90 minutes post-injection and a chest CT. Compared to lung uptake in the healthy volunteer, uptake was increased in IPF patients and enhanced in regions of reticulation and honeycomb patterns, consistent with localization of ^64^Cu-DOTA-ECL1i uptake in areas of active fibrotic remodeling. In most cases, radiotracer uptake was greatest in subpleural regions of the lung observed in sagittal sections and posterior coronal sections providing a spectrum of ^64^Cu-DOTA-ECL1i lung uptake (**Fig. 6A, B**). Whole-body images at 80-90 min post-injection showed less prominent uptake than the dynamic images due to rapid radiotracer clearance and metabolism, indicating that imaging during the first 60 minutes post-injection was superior for quantification.

**Fig. 6.**
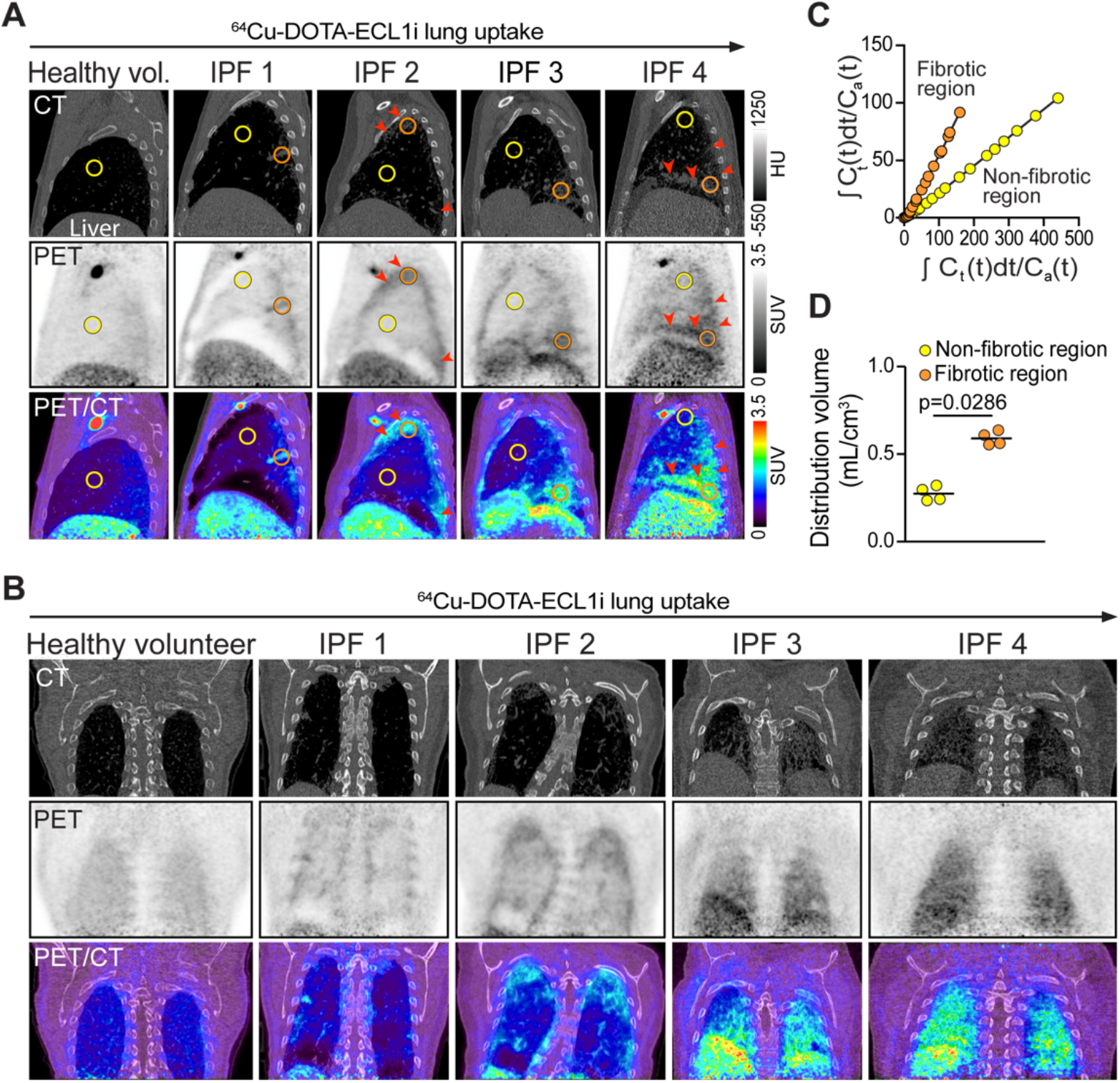
^64^Cu-DOTA-ECL1i PET/CT imaging in patients with IPF. A healthy volunteer and subjects with IPF (n=4) were injected with ^64^Cu-DOTA-ECL1i immediately prior to dynamic PET/CT imaging. (**A**) Representative sagittal plane CT, PET, and PET/CT-fused images showing subpleural uptake (arrows). Circled areas of non-fibrotic (yellow) and fibrotic (orange) lung were selected for Logan analysis. (**B**) Representative coronal plane images of lungs demonstrating subpleural uptake. (**C**) Logan plot demonstrates differing ^64^Cu-DOTA-ECL1i binding in regions of non-fibrotic and fibrotic tissue identified on PET/CT images from IPF2. (**D**) Differences in distribution volume of ^64^Cu-DOTA-ECL1i in non-fibrotic and fibrotic regions of IPF subjects (n=4). Significance determined by the Mann-Whitney U test. Images in (A) and (B) were arranged by increasing radiotracer uptake observed on PET.

**Table 1.** ^64^Cu-DOTA-ECL1i PET uptake metrics of subjects with IPF.

| Subject | Age (years) | Sex | O<sub>2</sub> use | FVC (%) | Fibrosis score* | ROI captured | Distribution volume | DV/ fibrosis ratio | SUV max | SUVmax fibrosis ratio |
| --- | --- | --- | --- | --- | --- | --- | --- | --- | --- | --- |
| IPF1 | 70 | M | No | 102 | 108 | Non-fibrotic | 0.25 | 1.96 | 1.20 | 1.63 |
|  |  |  |  |  |  | Fibrosis | 0.49 |  | 1.95 |  |
| IPF2 | 57 | F | No | 59 | 145 | Non-fibrotic | 0.22 | 2.63 | 0.75 | 2.64 |
|  |  |  |  |  |  | Fibrosis | 0.58 |  | 1.98 |  |
| IPF3 | 75 | M | No | 92 | 131 | Non-fibrotic | 0.34 | 1.62 | 1.50 | 2.13 |
|  |  |  |  |  |  | Fibrosis | 0.55 |  | 3.20 |  |
| IPF4 | 62 | M | Yes | 71 | 148 | Non-fibrotic | 0.33 | 2.18 | 2.70 | 2.20 |
|  |  |  |  |  |  | Fibrosis | 0.72 |  | 5.93 |  |

Abbreviations: FVC (%), Percent of predicted forced vital capacity; ROI, Region of interest; DV, Distribution volume; SUVmax, Maximum standard uptake value.
\*Fibrosis score measured as described (46).

^64^Cu-DOTA-ECL1i PET lung scans from the subjects were analyzed for standardized uptake value (SUV) and maximum SUV (SUVmax) (**Table 1**). Logan plots were used to determine the distribution volume (DV) using parametric maps derived from the dynamic images (*47*) and compared to that in healthy lungs (**Fig. 6C, Fig. S8**). The pulmonary artery was used as the blood reference region and high-uptake regions from sagittal and transverse images that best isolated vascular from parenchymal structures were selected for these assessments. Logan plots confirmed the increased uptake of radiotracer in areas of fibrosis, independent of blood flow. The average DV of the fibrotic regions (0.60 ± 0.04, n=4) was more than twice that of non-fibrotic regions (0.28 ± 0.04, n=4, p<0.05) (**Fig. 6D**). The SUVmax in these same fibrotic regions also tracked with distribution volume, validating the use of SUV. Metabolite analyses of blood samples obtained at 2-3 hours post-injection identified the major radioactive species as intact tracer (20-40%), free ^64^Cu (55-75%), which is not taken up in lungs (*48*), and negligible ^64^Cu-associated proteins (<5%). Thus, metabolites were unlikely to interfere with interpreting ^64^Cu-DOTA-ECL1i uptake as a measure of CCR2 expression. By these established measures of lung activity, two subjects had notably high uptake (IPF3, IPF4) compared to the two others (IPF1, IPF2). A subject who required oxygen supplementation (IPF4) had the highest ^64^Cu-DOTA-ECL1i uptake when compared to areas of similarly severe fibrosis identified in the other subjects (**Table 1**). The subject with the highest ratio of SUV or DV to fibrosis score (IPF2) developed rapid progression of disease, leading to lung transplantation. Moreover, assessment of coronal images showed differences in overall PET uptake between different subjects (**Fig. 6B**). This small number of patients was insufficient to define radiotracer uptake, fibrosis, and lung function relationships. However, these findings suggest that ^64^Cu-DOTA-ECL1i uptake may represent and important determinant of disease activity.

## Discussion

The nature of lung injury in IPF is not well defined and may be triggered by cell death or senescence (*1, 19, 37*). Nevertheless, for over two decades, the inflammatory response associated with fibrosis has been linked to bone marrow-derived CCR2^+^ lung monocytes and interstitial macrophages in animal models (*11–13, 49*). Most recently, analysis of lung cell from subjects with IPF using single cell transcriptional profiling has placed the interstitial macrophage in the profibrotic “niche”, serving as a driver for pulmonary fibrosis (*10, 15, 16, 21, 45*). Here, we complement those studies by showing a consistent, temporal, and spatial relationship between CCR2^+^ interstitial monocytes/macrophages and fibrotic regions using preclinical models and human tissue. We then extend those findings by showing the feasibility of a non-invasive PET imaging strategy for detection of CCR2^+^ lung cells during inflammation in the same preclinical models. Finally, we advance the potential to detect CCR2^+^ lung cells in patients by demonstrating perifibrotic uptake in first-in-human ^64^Cu-DOTA-ECL1i PET imaging in subjects with IPF. Our observations introduce the possibility that the inflammatory profibrotic niche may be non-invasively imaged in human fibrotic lung disease. The findings lead us to propose that measurable activity detected by ^64^Cu-DOTA-ECL1i PET may be used as a biomarker of immune cell activity in IPF.

Consideration of CCR2^+^ cells as a marker of profibrotic disease activity in the lung was judged by cellular activity following bleomycin injury, imaged at high resolution using CCR2-EGFP and *Ccr2* in situ hybridization, which localized to the perifibrotic region (**Figs. 1A, 3D**). The pattern was most striking in human tissues where CCR2^+^ cells were organized in distinct cell infiltrates around regions of marked fibrosis (**Figs. 4A, S5**). We found that CCR2^+^ interstitial macrophages in our model expressed the profibrotic genes, including *Cx3cr1*, identified by others in mouse models and human fibrotic lungs (*10, 14–16, 21, 45*) (**Figs. 1G, S3**). Our localization of the interstitial macrophages was consistent to photomicrographs in prior reports; Cx3cr1-expressing cells were directly adjacent to fibroblasts in the bleomycin model (*14*) and MERTK^+^ interstitial macrophages surround fibroblastic foci in human lung tissues (*21*). By scanning large regions of CCR2-immunostained tissue in human tissues, we could identify particularly dense CCR2^+^ infiltrates surrounding areas of honeycomb and scar, often in a “penumbra”-like pattern, suggesting a leading edge of CCR2^+^ cells driving the fibrotic process.

^64^Cu-DOTA-ECL1i uptake in tissue by autoradiography was also highly correlated with regions of CCR2 immunostaining, providing a level of validation for imaging of human subjects (**Figs. 4F, S7**). Ultimately, by using PET/CT, we found that patients with IPF had the most prominent signal in the subpleural regions. In limited studies, we linked the abundance of CCR2^+^ cells in regions of tissue from explanted lungs to the pre-transplant CT scans of those same regions (**Figs. 4A, S5**). However, in our small cohort of IPF subjects who underwent PET imaging, uptake was sometimes observed in regions without radiographic fibrosis. The signal in non-fibrotic lung may mark tissue in jeopardy of remodeling, hinting that ^64^Cu-DOTA-ECL1i PET could be a clinical tool for risk assessment (**Fig. 6**). We plan more rigorous validation of ^64^Cu-DOTA-ECL1i PET in patients with IPF undergoing lung transplantation, using explanted fibrotic lungs for comprehensive analysis of CCR2 levels and gene expression in specific anatomic locations, as guided by PET/CT.

It was essential we demonstrate that ^64^Cu-DOTA-ECL1i PET could detect changes in CCR2^+^ populations in treatment models. Modulation of CCR2 accumulation using IL-1β blockade and pirfenidone each significantly decreased PET radiotracer uptake. Previously it was shown that anti-IL-1β antibody blockade or genetic deficiency interrupts lung CCL2, egress of CCR2^+^ cells, and fibrosis (*19, 37, 38*). Indeed, even our treatment later in the course of inflammation (day 10 post bleomycin) showed effective diminution of CCR2^+^ cells and fibrosis, accompanied by decreased ^64^Cu-DOTA-ECL1i PET uptake. The treatment raises consideration for the use of monoclonal antibody against human IL-1β (canakinumab), or the IL-1R antagonist, anakinra, in IPF. A more clinically relevant approach was the treatment of bleomycin-induced fibrosis using pirfenidone, which reduced lung fibrosis, as previously reported (*23, 41*). A parallel fall in the burden of *Ccr2*^+^ cells was accompanied by decreased ^64^Cu-DOTA-ECL1i PET uptake. Although studied for many years, the precise pharmacology of pirfenidone has not been defined. At least one study using bleomycin injury reported a decrease in lung macrophages, CCL2, and CCR2 (*23*). Interestingly, pirfenidone inhibited NLRP3 inflammasome activation and IL-1β production in endotoxin-mediated lung injury of mice (*50*) pointing to the inflammasome as one pharmacologic target. Regardless, we now identify a potential drug whose effects may be reported by PET imaging with our non-invasive radiotracer.

There are unique, comparative advantages of ^64^Cu-DOTA-ECL1i PET uptake as a biomarker of IPF disease activity. As noted, chest imaging by high resolution CT scan is the current standard for diagnosis and a predictor of survival (*5, 6*), but it cannot yet be used to target a specific class of drug therapy. ^18^F-Fluorodeoxyglucose (^18^F-FDG)-PET can distinguish patients with severe IPF (*51*). However, ^18^F-FDG uptake is a non-specific marker of inflammation without a distinct molecular target. Newer PET radiotracers that measure α_v_β_6_ integrin (*52*), cathepsin (*53*), and collagen synthesis (*54*) may have related molecular phenotypes to provide complementary or alternative approaches to ^64^Cu-DOTA-ECL1i PET. Another feature of ^64^Cu-DOTA-ECL1i is the selection of Cu-64 as the radiolabel, which provides high specific activity and has straightforward radiochemistry. The stability and half-life of ^64^Cu (t_1/2_=12.7h) makes possible shipping of ^64^Cu-DOTA-ECL1i for future multicenter trials.

This study has limitations. First, therapeutic studies in mice are restricted to a few established experimental models with well-characterized trajectories and endpoints (*30*). Like most mouse models, bleomycin-induced fibrosis does not recapitulate all features of human disease, however, similar monocyte and macrophage populations are present after bleomycin and in IPF (*14, 15*). Second, CCR2^+^ monocytes/macrophages are but one inflammatory cell type identified in pulmonary fibrosis, and consideration for therapeutic depletion of this population may leave another pathogenic pro-fibrotic cell exposed to drive fibrosis or reveal a role of CCR2^+^ cells to resolve fibrosis (*55*). Third, as a first-in-human use of the radiotracer, a restricted number of subjects have been studied. We currently lack sufficient numbers of patients to draw firm conclusions related to the spectrum of lung uptake or relationship to therapy. We estimate that a sample size of 40 subjects with IPF would be required to achieve a 90% power to detect a significant correlation between ^64^Cu-DOTA-ECL1i uptake and whole-lung fibrosis score. Finally, although we have suggested that the effects of pirfenidone may be monitored by ^64^Cu-DOTA-ECL1i PET, it is possible that we will not observe the same effects in patients diagnosed late in the course of disease.

We conclude that detailed cellular tracking and lung localization of CCR2^+^ cells during the profibrotic processes expands knowledge of the role of these cells in pulmonary fibrosis, while development of a cell-targeted imaging probe may enhance clinical assessment of patients with IPF. The availability of ^64^Cu-DOTA-ECL1i PET as a cell-specific biomarker raises the distinct possibility that patients with IPF or other fibrotic diseases could be identified by quantitative CCR2^+^ measurement for specific drug treatment choices. Future studies may determine if ^64^Cu-DOTA-ECL1i PET can monitor pirfenidone therapy. At this time, it is unknown how the other approved anti-fibrotic agent, nintedanib, impacts the accumulation of CCR2^+^ cells or ^64^Cu-DOTA-ECL1i PET uptake. However, advancing the development of one of the many CCR2 antagonists that have been used in clinical trials (*56, 57*) may be an additional direction for targeted therapy to control a high CCR2^+^ cell condition. Future studies of ^64^Cu-DOTA-ECL1i PET as a broader tool for assessing inflammatory cell activity, and adapting it for fibrosis in other organs, such as the liver, which also share high CCR2^+^ cell populations (*58*).

## Materials and Methods

### Study design

The objective of these studies was to demonstrate the feasibility of the PET radiotracer ^64^Cu-DOTA-ECL1i to detect CCR2^+^ cells in patients with pulmonary fibrosis as a means to follow disease activity and for cell-targeted drug therapy. CCR2^+^ monocytes and interstitial macrophages were chosen as a target by reason of the pathologic roles in fibrosis of CCR2^+^ monocytes and lung interstitial macrophages in pulmonary fibrosis (*10, 14–20*). Complementary in vivo mouse and ex vivo human studies were designed with the goal of translating ^64^Cu-DOTA-ECL1i PET imaging to humans. The CCR2^+^ cell populations were localized relative to fibrotic regions in mouse lung tissues and characterized using a CCR2-reporter strain, single cell mass cytometry, *Ccr2* RNA in situ hybridization, and single cell transcriptomics (see Supplementary Methods), in parallel with ^64^Cu-DOTA-ECL1i PET uptake. Two mouse fibrosis models were used that are CCR2-dependent: the established bleomycin-induced fibrosis model and chronic radiation lung injury (*13, 36*). To determine if the ^64^Cu-DOTA-ECL1i PET uptake indicated modulation of CCR2^+^ monocytes and macrophage populations, therapeutic interventions were performed in the bleomycin injury model at the approximate onset of the fibrotic process, on day 10, and continued through 28, as recommended by a panel of experts (*40*). To modulate the proinflammatory effects of interstitial macrophages, both IL-1β blocking antibody and a clinically relevant medication, pirfenidone, were used. In each intervention, control and treated groups were assessed for fibrosis, CCR2^+^ cell abundance and location, imaged by ^64^Cu-DOTA-ECL1i PET. All animal studies were approved by the Institutional Animal Care and Use Committee at Washington University.

Ex vivo studies in human lung tissues explanted from subjects with IPF undergoing lung transplantation were designed to support the validity of in vivo imaging of subjects with IPF, and to localize CCR2^+^ cells relative to fibrotic regions. Donor lungs obtained from subjects without fibrosis that were unsuited for transplantation were used as a control. CCR2^+^ cells detected by immunofluorescence were associated with regions of fibrosis and correlation with the uptake of ^64^Cu-DOTA-ECL1i as detected by autoradiography in serial tissue samples. The primary objectives of the human in vivo studies were to determine dosimetry and safety of ^64^Cu-DOTA-ECL1i for PET imaging and detect uptake of radiotracer in the lung. ^64^Cu-DOTA-ECL1i was prepared for human imaging as described (*24*), under exploratory investigational new drug permission, using good manufacturing practices (see Supplementary Materials). Healthy volunteers included active cigarette smokers and never-smokers; all had normal lung spirometry testing and no diseases or medication use. Subjects were diagnosed with IPF by clinicians at the Washington University Interstitial Lung Disease Clinic, according to consensus clinical criteria (*5, 6*). The uptake of ^64^Cu-DOTA-ECL1i in lungs was measured in PET images using the method of Logan (*47*) and compared to clinical measures of lung function and a fibrosis score (*46*). Approval for human studies was provided by the Institutional Review Board at Washington University (IRB# 201606004), the Institutional Radioactive Drug Research Committee (protocol 826L) and the Food and Drug Administration (IND 137620). Written consent was obtained from all study participants.

### Mouse models

For the bleomycin-induced fibrosis model, male and female C57BL/6J wild-type mice (age, 8-10 weeks; approximately 25 grams; Jackson Labs) or mice with an EGFP-encoding DNA fragment inserted at the translation start site of *Ccr2* (B6C-Ccr2^tm1.1Cln^/J, #027619; Jackson Labs; referred to as CCR2^GFP/+^) were administered a single dose of bleomycin (Sigma), 3 U/kg, intranasal. For the single cell RNA sequencing studies, bleomycin oropharyngeal aspiration of 2 U/kg, was used. Co-housed, naïve mice served as controls to avoid confusing inflammatory responses induced by airway administration of saline vehicle. Weights were obtained every 1 to 3 days. Some mice given bleomycin were treated with isotype matched, polyclonal Armenian hamster IgG or anti-IL-1β (both from Bio X Cell), injected intraperitoneally, 200 μg, three times per week (*59*). Other cohorts were treated with pirfenidone (eEnovation Chemicals, D404655) mixed with rodent chow at 0.5% by weight, available ad libitum, as described (*43*). Lung radiation injury leading to fibrosis was induced in CD-1 mice (female, 5-7 weeks, 22-24 grams; Charles Rivers). A focal region of the right lung was irradiated with 20 Gy (*60*) delivered by the Small Animal Radiation Research Platform (Xstrahl).

### Mouse PET/CT and image analysis

Dynamic PET and CT scans were acquired using Inveon microPET/CT (Siemens) or Focus 220 PET (Concorde Microsystems) scanners, at 45 through 60 minutes following tail vein injection of ^64^Cu-DOTA-ECL1i (3.7 MBq per mouse). PET images were corrected for attenuation, scatter, normalization, and camera dead time, and co-registered with CT images. Both PET scanners were routinely cross-calibrated. The PET images were reconstructed with the maximum a posteriori algorithm and the organ uptake calculated as percent injected dose per gram (%ID/g) of tissue in three-dimensional regions of interest (ROIs) without correction for partial volume effect, using Inveon Research Workplace software (Siemens). Competitive PET blocking studies were performed with co-injection of non-radiolabeled ECL1i and ^64^Cu-DOTA-ECL1i at a molar ratio of 500:1 prior to imaging.

### Human PET/CT, imaging and dosimetry

(see Supplementary methods for human protocol and dosimetry details). Healthy volunteers received intravenous injection of 185-370 MBq of ^64^Cu-DOTA-ECL1i followed by four PET/CT scans (Biograph 40 PET/CT, Siemens) at 1 to 42 h. Images were acquired from the top of the skull through mid-thigh. Subjects with pulmonary fibrosis underwent a dynamic (0-60 minutes) PET scan of the thorax, which commenced upon intravenous injection of 296-370 MBq of ^64^Cu-DOTA-ECL1i and was followed by whole-body PET/CT imaging at 80-90 minutes post-injection. Dosimetry was calculated as previously described (*61*). Briefly, regions of interest (ROIs) were placed over the major organs using the CT as a guide. The time-activity curves were determined using all scans obtained to measure organ residence times. These data, plus the gamma radiation activity of urine collected after tracer injection, were used to calculate the dosimetry with OLINDA/EXM software (*62*). Organ SUV at each time point was determined directly from the images.

### Statistical analysis

Statistical analysis was performed using GraphPad Prism (Version 6.07). Differences between two groups were compared using the Mann-Whitney U test. Multiple medians were compared using the Kruskal-Wallis test followed by Dunn’s Multiple Comparison. Paired comparisons were analyzed using the Wilcoxon Signed Rank test. The Spearman correlation was used for analysis of ex vivo human tissue samples. P <0.05 was indicative of a statistically significant difference. Box plots show the median, a box representing first and third quartiles, and whiskers at 5^th^ and 95^th^ percentile, unless otherwise stated.

## Acknowledgments

We thank the study subjects for participation, and Luigi Adamo and Douglas Mann for assistance with pirfenidone dosing.

## Data and materials available

Single cell sequencing data will be deposited in NCBI GEO upon publication.

