## Supplemental methods, figures and tables for "Chemokine receptor 2-targeted molecular imaging in pulmonary fibrosis"

**Supplementary Material include:**

**Supplementary Materials and Methods:**

Synthesis and labeling ECL1i
Autoradiograph
Histochemistry, immunochemistry and microscopy
RNA in situ hybridization
Image analysis of tissue sections
Single cell isolation for mass cytometry
Cell preparation for mass cytometry antibody labeling
Antibody cocktail labeling for mass cytometry
Antibody titration for mass cytometry
Mass cytometry data analysis
Single cell isolation for RNA sequencing
Analysis of DropSeq data
Dosimetry methods

**Human Subjects Protocol for PET/CT Imaging**

**Supplementary Figures:**

**Fig. S1.** Method for localization of CCR2+ cells relative to fibrosis in mouse lung.
**Fig. S2.** Percentage of $CCR2^+$ cells detected by mass cytometry of lungs of mice treated with bleomycin.
**Fig. S3.** Profibrotic genes that cluster with mouse lung CCR2+ interstitial macrophages after bleomycin delivery.
**Fig. S4.** Detection of 64Cu-DOTA-ECL1i PET/CT activity in radiation-induced lung fibrosis in mice.
**Fig. S5.** Localization of CCR2+ cells in lung fibrotic tissue.
**Fig. S6.** Figure S6. Mass cytometry of cells from lungs of non-fibrotic donors and subjects with pulmonary fibrosis.
**Fig. S7.** Correlation of CCR2+ cells with 64Cu-DOTA-ECL1i uptake in fibrotic lung tissue..
**Fig. S8.** Analysis of 64Cu-DOTA-ECL1i PET images to determine radiotracer binding in the lungs from subjects with IPF.

**Supplementary Tables:**

**Table S1.** Antibodies used for mass cytometry
**Table S2.** Subjects providing lung tissue samples
**Table S3.** Subjects enrolled for 64Cu-DOTA-ECL1i PET/CT studies
**Table S4.** Estimated human dosimetry

**Supplementary Materials and Methods**

**Synthesis and labeling ECL1i.** $^{64}$Cu-DOTA-ECL1i was produce using previously described radiochemistry (*24-27*). The ECL1i peptide (LGTFLKC) was synthesized from D-form amino acids by CPC Scientific. DOTA-ECL1i was prepared by conjugating 1,4,7,10-tetraazacyclododecane-1,4,7,10-tetraacetic acid (DOTA) to the cysteine residue of ECL1i, using established methods (*24*). The crude conjugate was purified by high performance liquid chromatography to reach 99% chemical purity and verified by mass spectrometry. The DOTA-ECL1i conjugate was radiolabeled with $^{64}CuCl_2$ as described (*24*). $^{64}$Cu-DOTA-ECL1i for human imaging was prepared under exploratory investigational new drug permission, in compliance with the good manufacturing practices in the Biologic Therapy Core at Washington University. The final product contained 185- 370 MBq of $^{64}$Cu-DOTA-ECL1i with specific activity of 24.8-85.1 MBq/μg.

**Autoradiography.** Paraffin embedded human lung tissues on glass slides were rehydrated and processed for antigen retrieval by boiling in citrate-based Unmasking Solution (pH 6, Vector Laboratories) then blocked with 1% bovine serum albumin in PBS containing 0.1% Tween-20. The tissues were incubated with 11.1 MBq/m of $^{64}$Cu-DOTA-ECL1i for 15 min at room temperature with shaking, rinsed 3 times with purified water (MilliQ) and exposed for 30 min to a phosphor screen (GE Typhoon FLA 9500 Variable Mode Laser Scanner) at 50-micron resolution. Images of lung tissue autoradiography and serial sections of CCR2 detected by immunofluorescent staining were aligned using ImageJ. Fields of low and high $^{64}$Cu-DOTA-ECL1i signal (2.5 x 2.5 mm) were selected for analysis. The mean pixel intensity of autoradiography and immunofluorescence was calculated for each field.

**Histochemistry, immunochemistry and microscopy.** Mouse lungs were perfused with 8 mL of PBS injected into the vasculature through the right atrium, then inflated in situ with tissue fixative (Histochoice, Sigma) at 25 $cmH_2O$. The trachea was tied closed, lungs resected, and submerged in 10 mL of fixative for 24 h at room temperature. The tissue was paraffin embedded, and sectioned (5 μm) onto glass slides. To visualize collagen, tissue sections were stained by Gomori trichrome. For immunochemistry, deparaffinized and rehydrated tissue sections were heated in antigen unmasking solution pH 6.0 (Vector Laboratories) in a pressure cooker (Biocare Medical) for 10 min then cooled 30 min. The polyclonal anti-GFP antibody (Rockland) was used in mouse tissue and detected by using avidin-biotin amplification (Vectastain Elite ABC) and horseradish peroxidase substrate 3, 3-diaminobenzidine (DAB). Tissues were counterstained with hematoxylin. In human tissue, a monoclonal anti-CCR2 antibody (7A7; Abcam) was detected using tyramide amplification (Opal IHC Kit, PerkinElmer) and an Alexa Fluor 488-labeled secondary antibody (Thermo Fisher). DNA was stained with 4', 6 diamidino-2-phenylindole (DAPI) in mounting medium (Antifade, Sigma). Whole-slide images of stained tissue were captured using the NanoZoomer Whole-Slide Imaging System (Hamamatsu).

**RNA in situ hybridization.** *Ccr2* mRNA was visualized and quantified in paraffin-embedded mouse lung sections using the manufacturer protocol for RNAscope® fluorescent in situ hybridization (RNAscope® Multiplex Fluorescent v2 Assay kit, Advanced Cell Diagnostics). Briefly, sections were baked for 1 h at 60º C then deparaffinized. Target retrieval was performed by boiling sections for 15 min in a pressure cooker, followed by washing and treatment with protease for 30 min at 40º C. Tissues were incubated with a probe for CCR2 mRNA (Cat. Number 501681, Advanced Cell Diagnostic) for 2 h at 40º C and amplification was performed with RNAscope® amplifiers, horseradish peroxidase, and Opal 570 fluorophores (Perkin-Elmer). The sections were counterstained with DAPI and imaged with the Nanozoomer slide scanner.

**Image analysis of tissue sections.** The NDPITools plug-in for ImageJ was used to extract TIFF files from whole-slide images. For each tissue sample, two images were imported as separate layers into Photoshop (Adobe): (1) a trichrome-stained section and (2) a corresponding, immunohistochemical or immunofluorescent stained, serial section (**Fig. S1**). The images were cropped and the opacity of one layer adjusted such that both images could be viewed simultaneously. The images were aligned, and, using the "slice" tool, a grid of fields was created over the resulting composite (0.14 $mm^2$ for each field). The opacity was returned to 100% and the fields for both layers were exported as separate JPEG images using the "Save for Web" feature of Photoshop. Immunostained (DAB) cells in each field were counted by two independent observers using the "Cell Counter" function in ImageJ. Immunofluorescent, CCR2 signal was quantified in each field using the "MaxEntropy" setting of "Auto Threshold" in ImageJ. The DAPI signal was quantified using the “Default” setting of “Auto Threshold.” The severity of fibrosis in each corresponding field was scored using the “Analyse and Decide” function of the Blind Analysis Tool plug-in (ImageJ) by two independent observers using the modified Ashcroft scale (*31*).

**Single cell isolation for mass cytometry.** Human and mouse lung samples were handled in an identical manner except that the human lungs were not perfused. Mouse lungs were perfused with PBS and inflated with 1 mL digestion buffer containing 1.5 mg/mL Collagenase A (Roche), 80 μg/mL DNase I (Sigma), 5% fetal bovine serum, and 10 mM HEPES in PBS, injected into the trachea. The lungs were tied off, resected, and all lobes separated from the trachea and large airways. The mouse and human tissues were minced and submerged in digestion buffer and incubated at 37 ºC for 40 min with constant shaking (190 RPM) and gentle vortex mixing every 5 to 7 min. The digested tissue was dilute with 10 mL of ice cold PBS, mixed vigorously by vortex for 30 sec then passed through a 70-μm sieve (Falcon) to create a single cell suspension. Cells were centrifuged for 8 min at 500*g* at 4 ºC and collected. For the mouse tissues only, the red blood cells were then lysed by resuspending the pellet in 3 mL ACK Lysing Buffer (Lonza) which was neutralized with 20 mL ice-cold PBS and the cells collected by centrifugation. Human and mouse cells were suspended in FACS buffer (1% bovine serum albumin in PBS) and counted with a hemocytometer using trypan blue exclusion. Cells were suspended in freezing media (50% fetal bovine serum, 40% RPMI 1640, 10% DMSO) and cryopreserved in liquid nitrogen vapor until analysis by mass cytometry.

**Cell preparation for mass cytometry antibody labeling.** Each frozen aliquot of mouse or human cells was quickly thawed at 37 ºC and transferred to 10 mL of RPMI Plus media (RPMI 1640, 10% FBS, 10 U/mL heparin, 25 U/mL benzonase). Cells were pelleted at 300*g* for 10 min at room temperature and the supernatant aspirated. The cells were resuspended in 1 mL CyTOF PBS (10X PBS; Rockland Immunochemicals) diluted to 1X in MaxPar Water (Fluidigm). Cell number and viability was determined by the trypan dye exclusion method with manual counting using a hemocytometer. Cells suspended in CyTOF PBS were labeled with 1.25 μL of 1 mM stock of Fluidigm Cell-ID Cisplatin, 1.25 μM final concentration, for 1 minute at room temperature. Staining was quenched after the incubation with the addition of 5 mL of Fluidigm MaxPar Cell

Staining Buffer (CSB). Cells were pelleted (300*g*, 10 min, RT), the supernatant aspirated, and cells resuspended as $1x10^6$ cells/50 µL ($20x10^6$ cells/mL) in CSB.

**Antibody cocktail labeling for mass cytometry.** All antibodies were purchased conjugated by the manufacturer (Fluidigm), except Siglec-F, which was purchased carrier free (BD Biosciences) and conjugated using a MaxPar X8 antibody labeling kit (Fluidigm) according to manufacturer instructions (**Supplementary Table 4**). The antibody cocktail composing the panel was diluted in CSB to the previously determined titrated concentration (see below). Each sample was labeled with 50 µL of the antibody cocktail. For these experiments, two separate staining cocktails were utilized for cell surface and intracellular (i.e., granzyme B in human, GFP in mouse) antigens. A cell suspension of 50 µL in a 5 mL round-bottom polypropylene tube was incubated with 50 µL of the cell surface antibody cocktail, 1 h at 4 ºC. The cells were then washed twice with 2 mL CSB, pelleted (500*g*, 5 min, 4 ºC) and resuspended in 1 mL of 2% paraformaldehyde (EM grade, Electron Microscopy Sciences, "Fix Buffer") diluted in CyTOF PBS and fixed for 20 min at room temperature. Following fixation, cells were washed with 3 mL CSB, pelleted, and the supernatant aspirated. Cells were incubated with 50 µL CSB and 50 µL of intracellular antibody cocktail in Fix Buffer and incubated for 1 h, at 4 ºC. Cells were washed and collected again. To discriminate dead cells from live or single nucleated cells from doublets, MaxPar Intercalator-IR (Fluidigm) was prepared in Fix Buffer to a final concentration of 41.7 nM (3000X dilution of 125 µM stock). One mL of solution was added to each tube. Cells were incubated in the dark at 4 ºC. The following day, cells were washed twice with 2 mL CSB, then counted. Cells were subsequently washed with 2 mL MaxPar Water. The final cell concentration was adjusted to $1x10^6$ cells/mL in MaxPar Water with EQ calibration beads (Fluidigm) and filtered into cell strainer cap tubes. Data was then acquired on a CyTOF instrument (Helios, Fluidigm).

**Antibody titration for mass cytometry.** All antibodies were titrated for use by initially utilizing dilutions of 5-10 non-overlapping molecular weight antibodies on non-experimental specimens avoiding +1 and +16 channels for each antibody. To avoid "spill over" from abundant proteins into molecular weight gates that are 1 or 2 values higher or lower than the actual isotopically pure metal. Manufacturer recommendations typically result in too high of a concentration for very abundant surface proteins on cells like alveolar macrophages that are being evaluated with high sensitivity antibodies, such as HLA-DR. In addition, over time the isotopic metals can become oxidized and the oxidized isotope is also detected by the mass spectrometer. This effect is referred to as +16 interference as the oxidized isotope has a molecular weight that is one oxygen greater than the initial isotope. In the study of macrophages, these titration steps are important as most manufacturers have titrated the antibodies to detect these surface markers on cells such as monocytes, which produce much lower quantities of many of the proteins of interest.

**Mass cytometry data analysis.** Manual gating was performed using CytoBank software using standard parameters with the focus primarily on monocyte/macrophage cell lineages. For mice, after gating out $CD45^{-}$, $CD3^{+}$, $CD19^{+}$, $B220^{+}$, $NK1.1^{+}$, and $Ly6G^{high}$ cells ($Lin^{-}$), the cell phenotypes were: alveolar macrophages, $Lin^{+}/SiglecF^{high}$; interstitial macrophages, $Lin^{+}/SiglecF^{-}/CD64^{+}$; dendritic cells, $Lin^{+}/SiglecF^{-}/CD64^{-}/CD11c^{high}/MHCII^{high}$; inflammatory monocytes $Lin^{+}/SiglecF^{-}/CD64^{-}/CD11C^{low}/MHCII^{low}/Ly6C^{high}$; and patrolling monocytes, $Lin^{+}/SiglecF^{-}/CD64^{-}/CD11c^{low}/MHCII^{low}/Ly6C^{-}/F4/80^{+}$. Using this gating scheme, some $Ly6C^{+}$ interstitial macrophages remain that previously have been termed monocyte/macrophage (*15, 34*). For the purposes of this study, all $CD64^{+}$ cells are macrophages. For human cells, after gating out $CD45^{-}/CD3^{+}/CD19^{+}/CD56^{+}/CD66^{+}/Siglec8^{+}$ ($Lin^{-}$), the cell phenotypes were: alveolar macrophages, $Lin^{+}/HLA\text{-}DR^{+}/CD40^{+}/CD169^{+}$; interstitial macrophages, $Lin^{+}/HLA\text{-}DR^{+}/CD40^{+}/CD169^{-}$; dendritic cells, $Lin^{+}/CD80^{+}/CD40^{+}/CD14^{-}/CD16^{-}$; inflammatory monocytes, $Lin^{+}/CD40^{-}/HLA\text{-}DR^{low}/CD14^{+}/CD16^{-}$; and patrolling monocytes, $Lin^{+}/CD40^{-}/HLA\text{-}DR^{low}/CD16^{+}/CD14^{-}$.

**Single cell isolation for RNA sequencing.** Whole lungs from bleomycin treated and control mice were dissociated using a combination of mechanical and enzymatic dissociation. Briefly, lungs were rinsed with PBS and placed in digestion buffer including collagenase (4 mg/mL) and DNase (80 U/mL) in a GentleMacs dissociator using a C tube and the ‘lung1’ program (Miltenyi Biotec). The digest was continued at 37 °C for 30 min on a rotary shaker, followed by additional mechanical dissociation with the GentleMacs ‘lung2’ program. Cells were filtered through a 70-μm mesh. To clear the suspension of dead cells, red blood cells, and degraded nucleotides, the cell suspension was applied to an Optiprep (Sigma) gradient with 12%, 18%, and 30% layers and centrifuged at 600*g* for 15 min. Cells were extracted then washed twice with 50 mL 0.01% BSA in PBS, and pelleted at 1000*g* for 15 min. Cells were resuspended to a concentration of 100 cells/mL, and coencapsulated with barcoded beads. Prior to co-encapsulation, barcoded beads (ChemGenes) were diluted to 120 beads/mL. Droplets of 1 nL in volume were generated using microfluidic polydimethylsiloxane (PDMS) co-flow devices (FlowJEM Drop-seq chips) (*63*). Droplets were collected in a 50 mL RNase-free conical tube for a total run time of about 15 min. The collected emulsion was broken promptly with per-fluorooctanol, after which barcoded beads with captured transcriptomes were washed and centrifuged at 4 °C. RNA was reverse transcribed and exonuclease-treated using commercial kits. A total of 8000 beads per tube were used for PCR, amplified for 4+9 PCR cycles. PCR products were purified by addition of 0.6x Agencourt AMPureXP beads (Beckman Coulter #A63881). cDNA products meeting quality standards (>1000 bp average insertion size) were prepared for sequencing. cDNA from an estimated 5,000 cells were prepared and tagmented by Nextera XT (Illumina) using 600 pg of cDNA input. cDNA libraries were amplified (12 cycles) using custom primers as described (*64*). Amplified libraries were purified with a 0.6x volume of AMPure XP beads and quality was measured by Bioanalyzer. Libraries with an average length of 500-700 bp were submitted to Genome Technology Access Center of Washington University and sequenced on HiSeq 2500 and NovaSeq 6000 (Illumina).

**Analysis of DropSeq data.** Processing of paired-end sequencing reads to digital gene expression matrices with unique molecular identifier (UMI) counts was performed using STAR version 2.5.3a. Reads were aligned to the mm10 reference genome. Barcodes with less than 100 detected genes or >10% mitochondrial transcript counts were excluded from further analysis, along with cells with UMI counts > 5000 (suggestive of doublets). Count matrices were further analyzed using the R package Seurat version 3. For integrated analysis of bleomycin and control datasets, count matrices from control and bleomycin-treated mouse lungs were merged using canonical correlation analysis (CCA) using Seurat's *RunCCA* function, followed by normalization and scaling to limit cell-to-cell variation using the *NormalizeData* and *ScaleData* functions. Highly variable genes were calculated and subjected to principle components analysis, with the first 20 principle components provided as input to the clustering and dimensional reduction algorithms *FindClusters* and *RunUMAP*. Marker genes for each cluster were identified using the *FindAllMarkers* function, and cell types assigned using manual curation. From the whole data set, monocyte/macrophage populations were isolated using *SubsetData*, then subjected to repeat clustering using essentially the same pipeline and parameters as above. Myeloid lineage cell types were assigned as previously described (*15, 34*).

**Dosimetry Methods**. Data are obtained from PET/CT imaging from four studies obtained over approximately 42 h following a single injection of $^{64}$Cu-DOTA-ECL1i. A low-dose attenuation whole-body CT scan (30 effective mAs) was obtained from the skull vertex to the mid-thighs. The attenuation CT scan was obtained at normal, end-expiration to minimizes misregistration artifact. The PET images are obtained from the skull vertex to the mid-thighs after the CT scan. Regions of interest (ROIs) are traced on liver, kidney, urinary bladder content, spleen, blood pool, heart wall, fat, muscle, and total body. Activity in the legs (any part of the body not in the field of view) is estimated from the activity concentration in background. Background activity was measured in the shoulders. Leg and unaccounted activity were determined as follows:

- Legs (%ID) = Background*(Body Weight (g) – Total body ROI) / Injected activity
- Unaccounted = 100% - Sum Excreted (%ID) – Total body ROI (%ID) – Legs (%ID)

Unaccounted activity was assumed to be excreted in urine. Bladder content activity was tallied to generate bladder accumulation curves. Time activity curves were fitted with multi-exponential models and integrated analytically to yield organ residence time as described (*61*).

- Remainder = Max Residence Time – (Total observed – Total excreted)

Residence times were entered in OLINDA/EXM v1.1 for the appropriate adult model (*62*).

**Human Subjects Protocol for PET/CT Imaging**

**PET/CT Imaging with $^{64}$Cu-DOTA-ECL1i.** Imaging studies are performed in Barnes-Jewish Hospital at Washington University Medical Center in the Center for Clinical Imaging Research (CCIR). Two intravenous (i.v.) catheters will be placed, one for tracer injection and one for blood draws. Imaging data will be acquired on a Siemens Biograph 40 TruePoint Tomography PET/CT and reconstructed using 3-dimensional ordered subset expectation maximization (OSEM) with the attenuation correction CT.

**1. Dosimetry Study in Healthy volunteers**

**Inclusion Criteria: Healthy volunteers and healthy smokers**

- Men or women 21 years of age or older who have never smoked or current smokers who smoked at least 10 cigarettes per day (1/2 pack) and have smoked at least 100 cigarettes (5 packs) over the past month.
- Screening FEV1> 70% of predicted for healthy volunteers and current smokers
- Capable of lying still and supine within the PET/CT scanner for ~1 hour and follow instructions for breathing protocol during the CT portion
- No illicit drug use or other inhaled drug use (including pharmacologic agents, recreational agents, or illicit drugs) within the past year
- No known history of cardiac, pulmonary, hepatic or renal disease or diabetes
- No history of claustrophobia or other preventing condition that has previously or would interfere with completion of protocol-specified imaging sessions
- Able to comprehend and willing to follow instructions for the study procedures as called for by the protocol
- BMI ≤ 35

**Exclusion Criteria: Healthy volunteers and healthy smokers**

- Currently enrolled in another study using an investigational drug
- Pregnancy (confirmed by urine pregnancy test)
- Active symptoms or history of cardiac, pulmonary, hepatic, or renal disease or diabetes
- Currently taking any prescription medications
- Presence of an implanted device that is incompatible with CT scanning
- Creatinine > 1.30 mg/dL, AST > 50 Units/L, ALT > 55 Units/L, or total bilirubin > 1.2 mg/dL

**Protocol for Dosimetry Study in Healthy Volunteers**

**Day 1: Screen visit**

1. The informed consent document will be reviewed again with the participant. The participant will sign the consent at this time.
2. Vital signs and pulse oximetry will be obtained, and height, weight, and BMI documented.
3. A urine pregnancy test will be conducted to confirm non-pregnancy (if applicable).
4. Baseline Complete Blood Count (CBC) and Complete Metabolic Panel (CMP) samples will be drawn and a urine sample collected.
5. Pulmonary function tests (spirometry only) and ECG will be performed.

**Day 2-4: PET/CT imaging visit**

This portion of visits will take up to 3 days for each participant. The visits will be scheduled within two weeks of the Screen Visit.

1. Participants will arrive at the CCIR.
2. Vital signs, pulse oximetry, height, weight, and BMI will be repeated.
3. A urine pregnancy test will be conducted to confirm non-pregnancy (if applicable).
4. PET/CT and with $^{64}$Cu-DOTA-ECL1i will be obtained as described in Section 4.2.1.
5. Vital signs will be re-checked upon completion of each PET/CT scan.
6. Another set of CBC and CMP blood samples will be obtained, and ECG will be performed after all scans have been completed. An additional urine sample will also be obtained for repeat urinalysis (see below).
7. The volunteer will be asked to urinate into a container for volume measurement immediately after completing the PET/CT scan session on the first day within 1 hour of tracer injection. After that, the volunteer will be asked to urinate before and after each scan session, as they are able. A urine sample from each void will be collected for gamma counting and used in the dosimetry calculation.

All participants will be called the day after completing all the imaging procedures to assess for any changes in health status.

#### 2. Lung Imaging in Subjects with ILD and lung fibrosis

Inclusion Criteria: Participants with ILD and lung fibrosis

- Men or women 21 years of age or older who have well documented lung disease with fibrosis followed at the Lung Center clinic or clinician diagnosis of radiation-induced fibrosis and/or radiation-induced pneumonitis at the Radiation Oncology clinic at Washington University
- Pulmonary function tests within the past one year demonstrating an FVC > 50% of predicted
- Glomerular filtration rate (GFR) >50 mL/min determined using the Chronic Kidney Disease Epidemiology Collaboration Equation (CDK-EPI)
- No active liver disease based on clinical evaluation
- No uncontrolled extra-pulmonary disease based on clinical evaluation
- Capable of lying still and supine within the PET/CT scanner for ~1 hour and follow instructions for breathing protocol during the CT portion
- No illicit drug use or smoking tobacco use (including recreational agents or illicit drugs) within the past year
- No history of claustrophobia or other preventing condition that has previously or would interfere with completion of protocol specified imaging sessions
- Able to comprehend and willing to follow instructions for the study procedures as called for by the protocol
- BMI ≤ 35

Exclusion Criteria: Participants with ILD and lung fibrosis

- Currently enrolled in another study using an investigational drug
- Pregnancy (confirmed by urine pregnancy test)
- Pulmonary disease requiring more than 4 liters/minute of supplemental oxygen at rest
- GFR ≤ 50 mL/min, AST > 50 Units/L, ALT > 55 Units/L, or total bilirubin > 1.2 mg/dL

- Presence of an implanted device that is incompatible with CT scanning

**Lung Imaging Study in Subjects with lung fibrosis.**

1. Participants will arrive at the CCIR. CBC and CMP blood samples and urinalysis will be obtained, and ECG performed.
2. Vital signs, pulse oximetry, height, weight, and BMI will be obtained
3. A urine pregnancy test will be conducted to confirm non-pregnancy (if applicable).
4. PET/CT imaging: A low-dose attenuation chest CT scan (30 effective mAs) will be obtained at normal end-expiration followed by a 60-minute dynamic PET image acquisition that starts at the time of $^{64}$Cu-DOTA-ECL1i injection (5 to 10 mCi) using the following framing schedule: 24 5-second, 3 1-minute, 5 3-minute, and 8 5-minute frames (40 frames total). Blood may be drawn during this time to determine the input function and assess radiotracer protein binding (approximately 45 mL total for this purpose).
5. After completing the dynamic scan, the participant will be allowed to stand up and use the restroom prior to the next acquisition. The second scan will then be obtained 1-3 hours later as follows. A low-dose attenuation whole-body CT scan (30 effective mAs) will be obtained from the skull vertex to the mid-thighs at normal end-expiration followed by PET images of the same regions.
6. Blood samples may be collected at the time of each scan to determine the presence of radiotracer metabolites by high performance liquid chromatography (HPLC) (approximately 15 mL in total) when staff are available.
7. Vital signs will be re-checked upon completion of each PET/CT scan.
8. Another set of CBC and CMP blood samples will be obtained, and ECG will be performed after all scans have been completed. An additional urine sample will also be obtained for repeat urinalysis (see below).
9. Vital signs, CBC and CMP blood tests urinalysis will be repeated at the conclusion of the imaging sessions. A total of approximately 65 mL of blood may be collected for these blood draws and the blood draws during the scan.
10. All participants will be called the day after completing all the imaging procedures to assess for any changes in health status.

**SUPPLEMENTARY FIGURES**

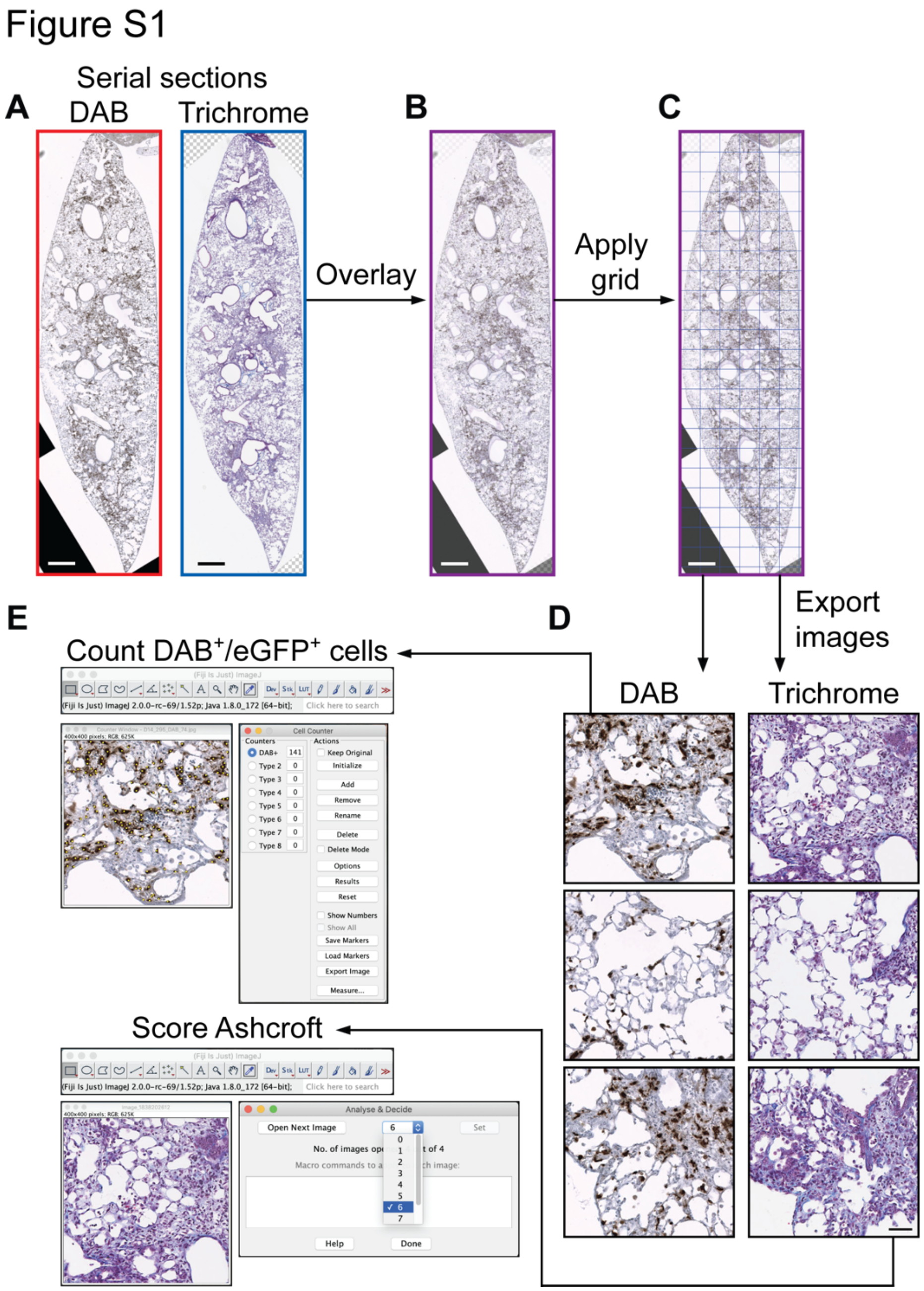

**Fig. S1. Method for localization of CCR2$^+$ cells relative to fibrosis in mouse lung.** C57BL/6 CCR2$^{GFP/+}$ mice were administered intranasal bleomycin and lungs assayed at day 14. All segments of mouse lungs were embedded in paraffin and tissue sections stained. Entire lung sections were imaged across the slide using the Nanozoomer slide scanner. (**A**) An image of mouse lung with CCR2$^{GFP/+}$ cells identified by an anti-GFP antibody and the chromogen 3, 3-diaminobenzidine (DAB, brown, left) and an image of a serial tissue section stained for collagen using trichrome (blue, right). (**B**) Images overlaid by adjusting global opacity in Photoshop. (**C**) A grid applied using the Photoshop "slice" tool to create fields that are 0.14 mm$^2$ each across the image. (**D**) Examples of fields exported by Photoshop from the DAB ($CCR2^{GFP/+}$ cells) and trichrome stained layers of the full image in C. (**E**) Fields were randomized and $CCR2^{GFP/+}$ cells were counted and fibrosis severity scored (by modified Ashcroft) using the "Analyse and Decide" and "Cell Counter" tools, respectively, in ImageJ. A similar system was used for analysis of lung tissue stained for CCR2 using immunofluorescence in mouse and human samples. A-C, Bar=500 mm; D, E Bar=50 μm.

Figure S2

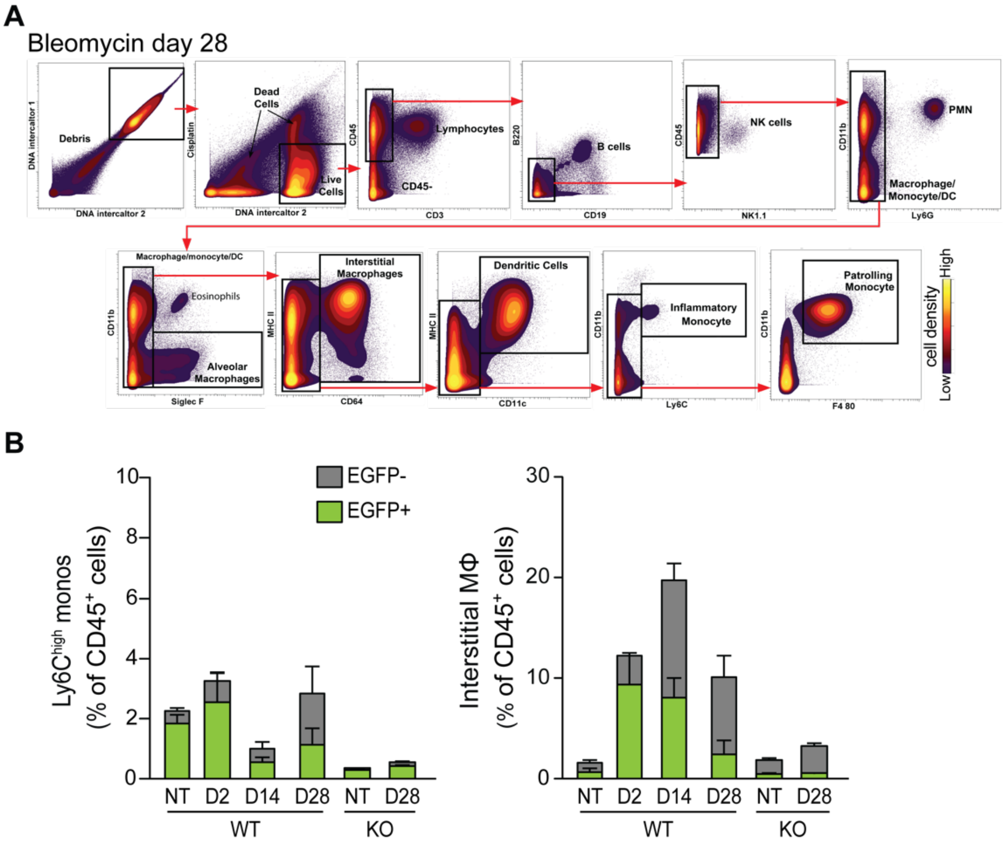

**Fig. S2. Percentage of CCR2$^{+}$ cells detected by mass cytometry of lungs of mice treated with bleomycin. (A)** Gating strategy for identification of immune cell populations. Single cells were isolated from the lungs of mice 28 days after administration of bleomycin. Myeloid cells are defined as follows: Ly6C$^{high}$ inflammatory monocytes (monos; SiglecF$^{-}$/CD64$^{-}$/Ly6C$^{high}$), interstitial macrophages (MΦ; SiglecF$^{-}$/CD64$^{+}$), dendritic cells (DCs; SiglecF$^{-}$/CD64$^{-}$/CD11c$^{high}$/MHCII$^{high}$), patrolling monocytes (monos; SiglecF$^{-}$/CD64$^{-}$/CD11c$^{low}$/MHCII$^{low}$/ Ly6C-/F4/80$^{+}$), and alveolar macrophages (MΦ; SiglecF$^{high}$). (**B**) Percentage of inflammatory monocytes and interstitial macrophages in the lungs of C57BL/6 CCR2$^{GFP/+}$ or CCR2 knockout (KO, CCR$^{GFP/GFP}$) littermates either non-treated (NT) or at timepoints after bleomycin injury (day 2, 14, 28). Shown are the median with interquartile range of percent of CD45$^{+}$ cells of each cell type (gray bars) and the percent that are CCR2$^{GFP/+}$ (green bars). n=3-5 mouse lungs/time points.

### Figure S3

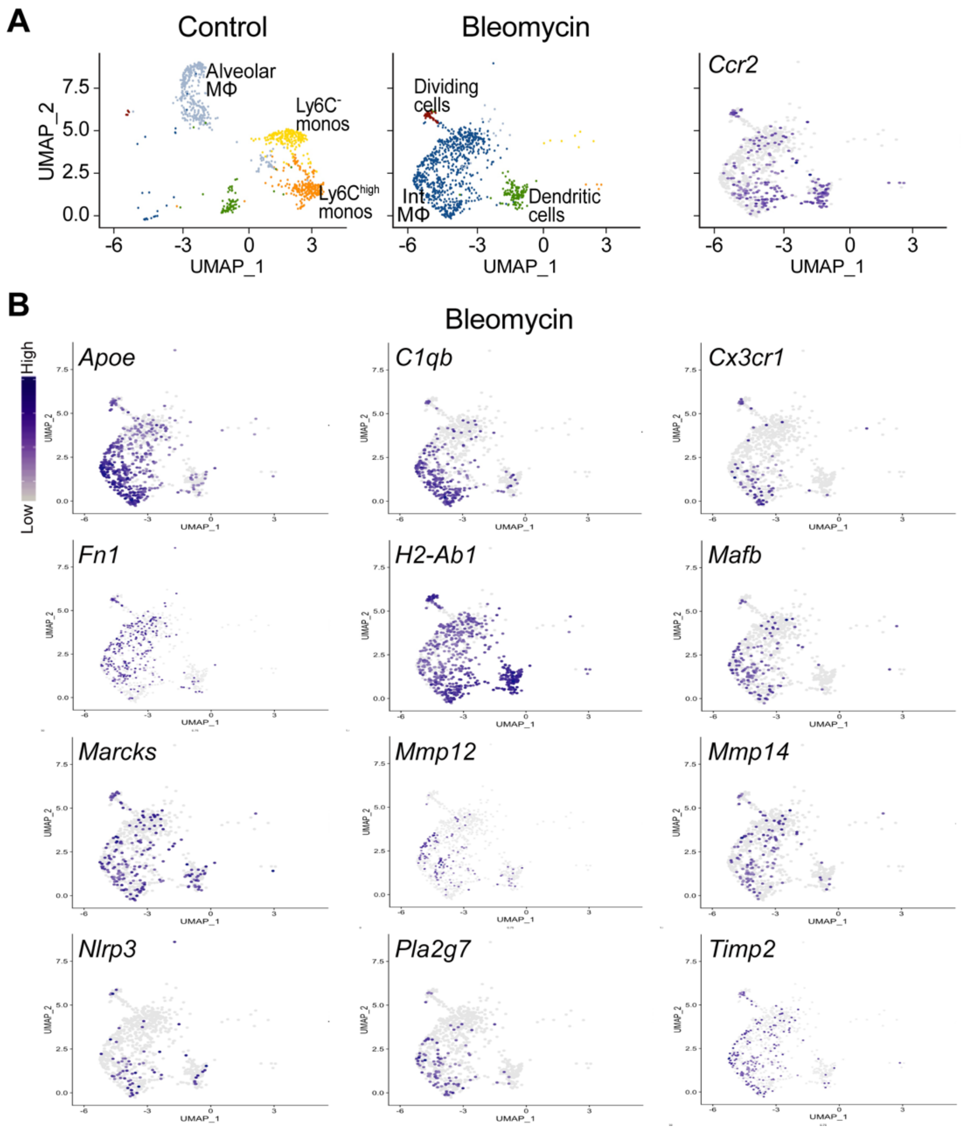

**Fig. S3. Profibrotic genes that cluster with mouse lung CCR2$^+$ interstitial macrophages after bleomycin delivery.** Mice were administered intratracheal control saline or bleomycin. Twenty days later, single cells were isolated and analyzed for transcription profiles. A total of 15,187 cells, with 533 genes and 853 unique molecular identifiers per cell were detected in control and bleomycin treated samples. Transcriptional profiles in cells were visualized by the Uniform Manifold Approximation and Projection (UMAP) dimensional reduction technique using Suratt software. (**A**) Representative differences in transcription of *Ccr2* in myeloid single cell populations isolated from lungs of wild-type mice. Cell types clustered by identity as shown in Fig 1. (**G**) Cells expressing *Ccr2* proinflammatory and profibrotic genes in interstitial macrophages post-bleomycin, in addition to those shown in Fig. 1. Shown are representative data from one mouse lung (n=2 mice/condition).

Figure S4

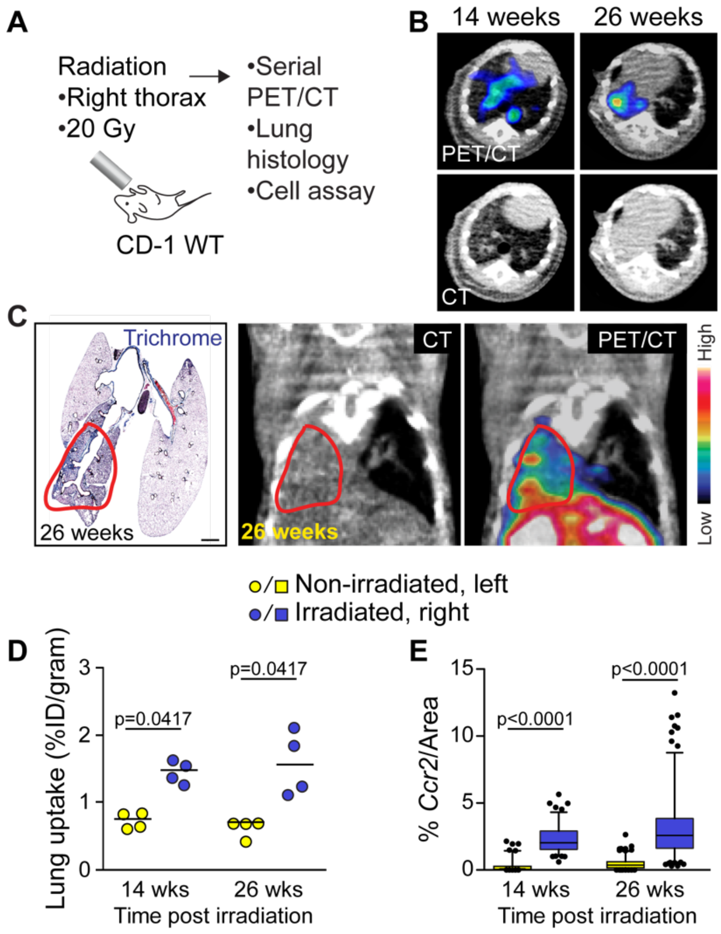

**Fig. S4. Detection of** $^{64}$**Cu-DOTA-ECL1i PET/CT activity in radiation-induced lung fibrosis in mice.** (**A**) The right thorax of mice was irradiated using 20 Gy. Mice were assayed 14 or 26 weeks later. (**B**) At the indicated time, mice were injected with $^{64}$Cu-DOTA-ECL1i, then underwent PET/CT. (**C**) Transverse section of mouse lungs stained with trichrome with companion PET image. (**D**) The medium lung uptake of irradiated, right and non-irradiated, left lungs (n=4 mice per timepoint). (**E**) Quantitation of *Ccr2* in the right and left lung using in situ hybridization in lung sections obtained at the indicated time post-irradiation (118-173 fields/lung, n=3 mice/time point). Significance was determined in D by the Wilcoxon Signed Rank test and in E, the Mann-Whitney U test. In C, Bar=1000 μm.

Figure S5

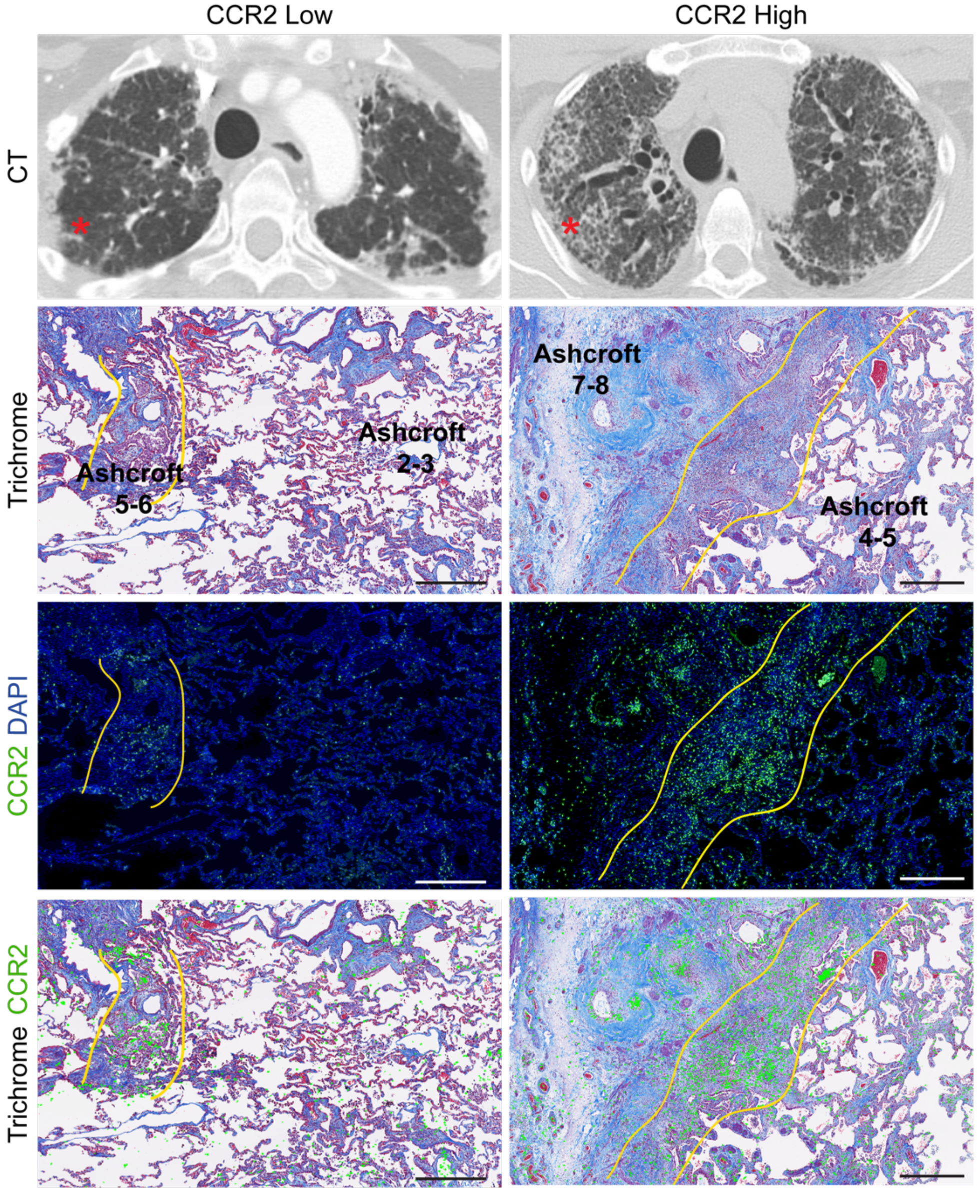

**Fig. S5. Localization of CCR2$^{+}$ cells in lung fibrotic tissue.** Chest computed tomography (CT) of subjects with end-stage pulmonary fibrosis obtained prior to undergoing lung transplantation. Explanted lungs were processed using an established biobank protocol to sample different regions. Representative examples are shown. Chest CT shows the region of lung sampled (asterisk). Serial sections of tissue were stained with trichrome and immunostained with anti-CCR2 antibody identified with Alexa Fluor 488 (green) and DAPI stained nuclei (blue). The bottom panels are an overlay. Regions of abundant CCR2$^{+}$ cells (outlined in yellow) relative to fibrosis are shown. Bar = 500 μm.

Figure S6

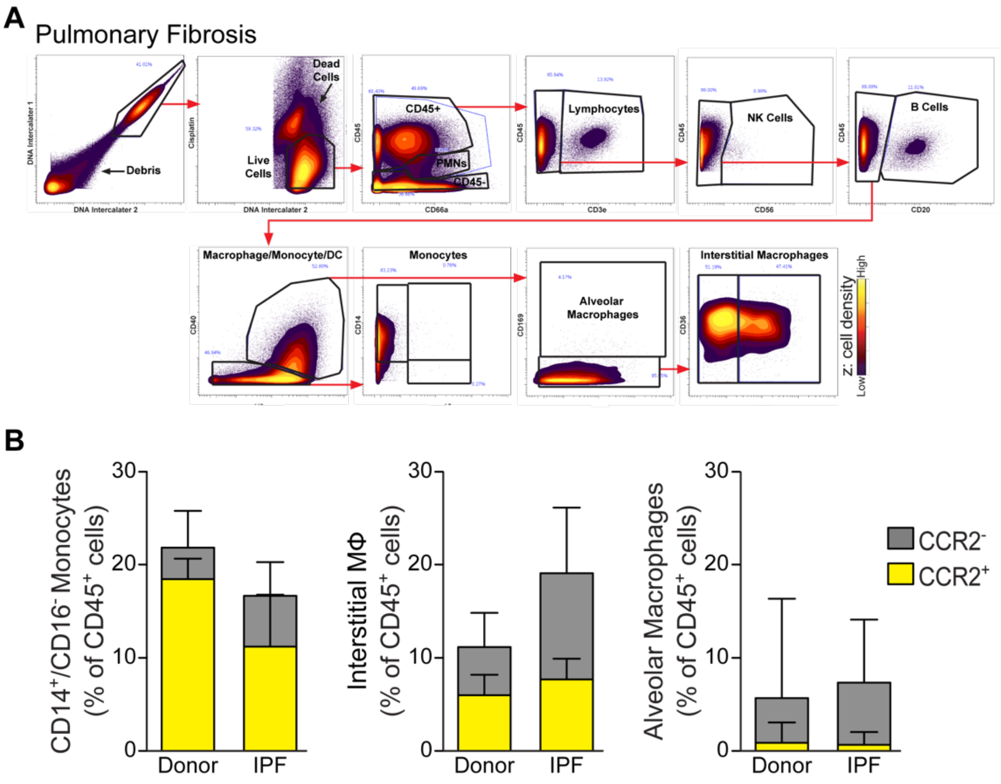

**Fig. S6. Mass cytometry of cells from lungs of non-fibrotic donors and subjects with pulmonary fibrosis.** Single cells were isolated from the lungs of non-fibrotic lungs donated, but not used for transplantation (Donor) and subjects with end-stage pulmonary fibrosis whose lungs were remove in the course of lung transplantation (PF). (**A**) Gating strategy for identification of immune cell populations in a subject with pulmonary fibrosis. The cell phenotypes were: alveolar macrophages, $Lin^+/HLA\text{-}DR^+/CD40^+/CD169^+$; interstitial macrophages, $Lin^+/HLA\text{-}DR^+/CD40^+/CD169^-$; dendritic cells, $Lin^+/CD80^+/CD40^+/CD14^-/CD16^-$; inflammatory monocytes, $Lin^+/CD40^-/HLA\text{-}DR^{low}/CD14^+/CD16^-$; and patrolling monocytes, $Lin^+/CD40^-/HLA\text{-}DR^{low}/CD16^+/CD14^-$. (**B**) Percent of $CD45^+$ cells that were interstitial monocytes, inflammatory macrophages, and alveolar macrophages. Shown are the median and interquartile range (n=6 donor, n=11 PF).

Figure S7

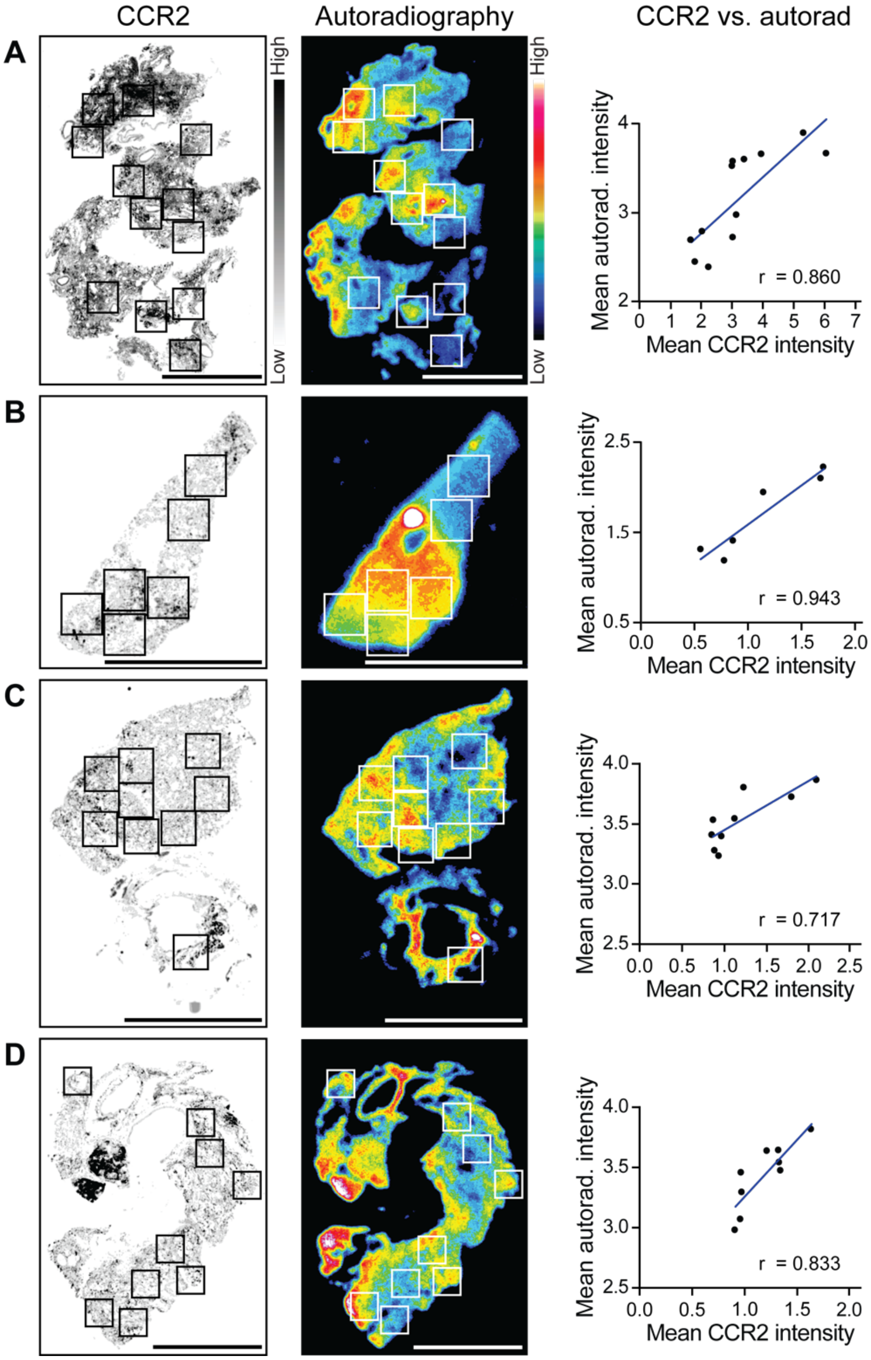

**Fig. S7. Correlation of CCR2$^{+}$ cells with $^{64}$Cu-DOTA-ECL1i uptake in fibrotic lung tissue.** Lung tissues were obtained from subjects with end-stage pulmonary fibrosis lungs undergoing lung transplantation. (**A-D**) Representative examples are shown. Serial lung tissue sections were immunostained for CCR2 (left) or incubated with $^{64}$Cu-DOTA-ECL1i followed by detection of binding by autoradiography (center). Entire tissue sections on a microscopy slides were scanned to create photomicrographs and the images aligned using the “Align Image by Line ROI” function in ImageJ. Areas of high and low CCR2 immunostaining were selected (boxes) and then transposed onto the corresponding area of the autoradiograph images. The pixel intensity in each box was measured for Spearman correlation (right). Bar=10 mm.

Figure S8

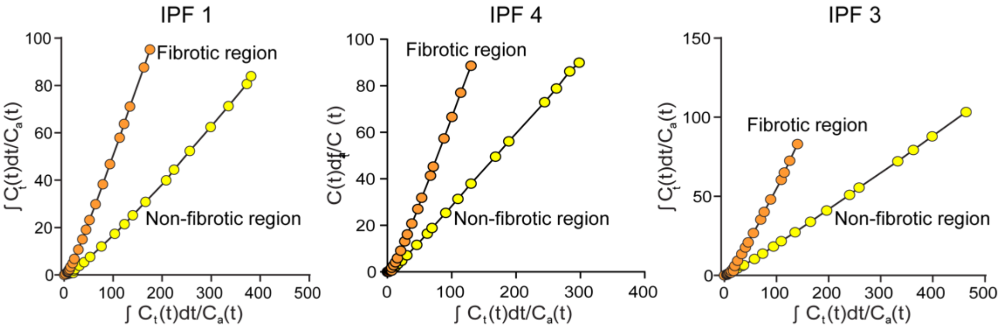

**Fig. S8. Analysis of $^{64}$Cu-DOTA-ECL1i PET images to determine radiotracer binding in the lungs from subjects with IPF.** Subjects with IPF were administered intravenous $^{64}$Cu-DOTA-ECL1i followed by PET/CT imaging. Regions of high and low fibrosis were selected for analysis as shown in Fig. 6 for calculation of specific $^{64}$Cu-DOTA-ECL1i binding.

**Table S1. Antibodies used for mass cytometry**

| Metal label | Human | | | | Mouse | | | |
|---|---|---|---|---|---|---|---|---|
| | Specificity | Clone | Titration | Catalog # | Specificity | Clone | Titration | Catalog # |
| **89Y** | CD45 | HI30 | 0.1 | 3089003B | CD45 | 30-F11 | 0.25 | 3089005B |
| **141Pr** | CD49d | 9F10 | 0.25 | 3141004B | Ly6G/C (Gr1) | RB6-8C5 | 0.25 | 3141005B |
| **142Nd** | CD19 | HIB19 | 0.1 | 3142001B | CD11c | N418 | 0.25 | 3142003B |
| **143Nd** | CD127 (IL17R) | A019D5 | 0.1 | 3143012B | | | | |
| **144Nd** | CD11b | ICRF44 | 0.5 | 3144001B | CD16/32 | 93 | 0.5 | 3144009B |
| **145Nd** | CD4 | RPA-T4 | 0.33 | 3145001B | CD69 | H1.2F3 | 0.25 | 3145005B |
| **146Nd** | CD64 | 10.1 | 0.25 | 3146006B | | | | |
| **147Sm** | CD20 | 2H7 | 1 | 3147001B | CD36 | No.72-1 | | 3147013B |
| **148Nd** | CD16 | 3G8 | 0.5 | 3148004B | CD11b(Mac1) | M1/70 | 0.25 | 3148003B |
| **149Sm** | CD66 | CD66a-B1.1 | 0.33 | 3149008B | CD19 | 6D5 | 0.25 | 3149002B |
| **150Nd** | CD223(Lag3) | 11C3C65 | 1 | 3150030B | Ly6c | HK1.4 | 0.25 | 3150010B |
| **151Eu** | CD123 | 6H6 | 0.5 | 3151001B | CD25 | 3C7 | 0.25 | 3151007B |
| **152Sm** | CD36 | 5-271 | 0.25 | 3152007B | CD3e | 145-2c11 | 1 | 3152004B |
| **153Eu** | CCR2 | K036C2 | 0.5 | 3153023B | CD274(PD-L1) | 10F.9G2 | 0.1 | 3153016B |
| **154Sm** | CD163 | GHI/61 | 0.5 | 3154007B | Ter-119 | TER119 | 0.25 | 3154005B |
| **155Gd** | CD279 (PD1) | EH12.2H7 | 0.5 | 3155009B | SiglecF | E50-2440 | 0.5 | BD552125 |
| **156Gd** | CD86 | IT2.2 | 0.25 | 3156008B | CD14 | Sa142 | 0.25 | 3156009B |
| **158Gd** | CD169 | 7-239 | 0.5 | 3158027B | | | | |
| **159Tb** | CD11c | Bu15 | 0.1 | 3159001B | F4/80 | BM8 | 0.5 | 3159009B |
| **160Gd** | CD14 | M5E2 | 0.5 | 3160001B | CD62L | MEL-14 | 0.25 | 3160008B |
| **161Dy** | CTLA-4 | 14D3 | 0.5 | 3161004B | | | | |
| **162Dy** | CD80 | B7-1 | 1 | 3162010B | CD1d | 1B1 | 0.1 | 3162020B |
| **163Dy** | CD172 | SE5A5 | 1 | 3163017B | | | | |
| **164Dy** | Siglec-8 | 7C9 | 0.5 | 3164017B | CX3CR1 | 8F1/CXCR1 | 0.25 | 3142009B |
| **165Ho** | CD40 | 5C3 | 0.5 | 3165005B | CD31(PECAM1) | 390 | 0.5 | 3165013B |
| **166Er** | CD44 | Bj18 | 0.5 | 3166001B | CD326(EpCAM) | G8.8 | 0.5 | 3166014B |
| **167Er** | CD27 | L128 | 0.5 | 3167006B | | | | |
| **168Er** | CD206 | 15-2 | 0.1 | 3168008B | CD8a | 536.7 | 0.25 | 3168003B |
| **169Tm** | CD45RA | HI100 | 1 | 3169008B | GFP | | 0.5 | |
| **170Er** | CD3e | UCHT1 | 0.25 | 3170001B | NK1.1 | PK136 | 0.5 | 3170002B |
| **171Yb** | CD68 | Y1/82A | 0.5 | 3171011B | CD44 | IM7 | 0.25 | 3171003B |
| **172Yb** | CD38 | HIT2 | 0.33 | 3172007B | CD4 | RM4-5 | 0.25 | 3172003B |
| **173Yb** | CXCR4 | 12G5 | 0.25 | 3173001B | | | | |
| **174Yb** | HLA-DR | L243 | 0.33 | 3174001B | MHC II (1A/1E) | M5/114.15.2 | 0.5 | 3174003B |
| **175Lu** | CD274 (PDL1) | 29E.2A3 | 1 | 3175017B | CD207 | 4C7 | 1 | 3175016B |
| **176Yb** | CD56 | NCAM16.2 | 0.1 | 3176008B | B220 | RA3-6B2 | 0.5 | 3176002B |
| **178Os** | Background | | | | Background | | | |
| **191Ir** | DNA1 | | | 201192B | DNA1 | | | 201192B |
| **193Ir** | DNA2 | | | 201192B | DNA2 | | | 201192B |
| **195Pt** | cisplatin | | | 201194 | cisplatin | | | 201194 |

**Table S2. Subjects providing lung tissue samples**

| Sample | Age (yrs) | Sex | Race | Health status | Cigarette Smoking | Assays |
|---|---|---|---|---|---|---|
| **Control donor non-fibrosis samples** | | | | | | |
| **D1** | 56 | F | W | Cardiovascular disease | 7.5 pack yrs | MC |
| **D2** | 30 | Unk | Unk | Unk | 6 pack yrs | MC |
| **D3** | 55 | F | Unk | Unk | Unk | MC |
| **D4** | 53 | F | W | Cardiovascular disease | 10 pk yrs | IF |
| **D5** | 19 | M | W | Blunt injury | None | IF |
| **D6** | 62 | M | W | Intracerebral hemorrhage | Unk | IF, TC |
| **D7** | 50 | M | W | Ischemic stroke | Unk | MC |
| **D8** | 51 | M | W | Cerebrovascular stroke | Unk | IF, MC |
| **D9** | 61 | F | W | Intracerebral hemorrhage | None | MC |
| **Pulmonary fibrosis samples** | | | | | | |
| **F1** | 70 | M | W | Familial IPF/UIP | 25 pack yrs, none 28 yrs | MC |
| **F2** | 63 | M | W | IPF with UIP | 26 pack yrs, none 32 yrs | MC |
| **F3** | 62 | M | W | Familial IPF/UIP | None | MC |
| **F4** | 69 | F | W | NSIP | None | TC, IF, AR |
| **F5** | 65 | F | W | NSIP | None | IF, MC |
| **F6** | 70 | M | A | IPF/UIP | None | CT, TC, IF |
| **F7** | 59 | M | W | UIP | None | TC, IF, AR |
| **F8** | 59 | F | W | IPF/UIP | None | IF |
| **F9** | 42 | F | W | Familial IPF/UIP | None | CT, TC, IF |
| **F10** | 66 | F | W | PF | 7 pack yrs, none 33 yrs | IF |
| **F11** | 69 | M | W | IPF/UIP | 112 pack yrs, none 3 yrs | IF |
| **F12** | 58 | M | W | NSIP | None | TC, IF, AR |
| **F13** | 64 | M | W | PF | 43 pack yrs, none 2 yrs | TC, IF, AR |
| **F14** | 65 | F | W | IPF with UIP | None | MC |
| **F15** | 57 | F | B | IPF | 15 pack yrs, at least 1 yr | MC |
| **F16** | 61 | F | W | PF | 0.5 PPD x many yrs, none 9 yrs | MC |
| **F17** | 58 | M | W | IPF/UIP | 0.75 PPD x many yrs, none 4 yrs | MC |
| **F18** | 53 | M | W | IPF/UIP | None | MC |
| **F19** | 66 | M | W | IPF/UIP | None | MC |
| **F20** | 75 | M | W | IPF/UIP | 60 pack yrs, none 30 yrs | MC |
| **F21** | 57 | F | W | Familial IPF/UIP | None | CT, TC, IF |

**Abbreviations:** Unk, unknown; F, female; M, male; A, Asian; B, black; W, white; IPF, Idiopathic pulmonary fibrosis; UIP, usual interstitial pneumonia; NSIP, non-specific interstitial pneumonia; PPD, packs of cigarettes per day; PF, Pulmonary fibrosis; CPFE; combined pulmonary fibrosis and emphysema; PPD, Unk, unknown; number of packs smoked per day; Yrs, years.

**Assays:** MC, mass cytometry; IF, immunofluorescence for CCR2; Autoradiography for $^{64}$Cu-DOTA-ECL1i ; CT, chest computed tomography

**Table S3. Subjects enrolled for $^{64}$Cu-DOTA-ECL1i PET studies**

| Subject | Age (yrs) | Sex | Race | Wt (kg) Ht (cm) BMI | Health status | Cigarette smoking (PPD or pack years) | Drug treatment | FVC (%) | Injected dose (mCi) | Comment |
|---|---|---|---|---|---|---|---|---|---|---|
| **Dosimetry and healthy control subjects** | | | | | | | | | | |
| **H1** | 23 | F | W | 54.7<br>167<br>19.6 | Healthy | None | None | >90 | 5.2 | |
| **H2** | 27 | M | W | 76.2<br>179<br>23.8 | Healthy | None | None | >90 | 5.6 | |
| **H3** | 28 | F | W | 54.7<br>163<br>20.6 | Healthy | 5 pack yrs; active | None | >90 | 9.2 | |
| **H4** | 37 | M | B | 94.3<br>185<br>27.7 | Healthy | None | None | >90 | 10.4 | |
| **H5** | 43 | F | W | 61.2<br>166<br>22.2 | Healthy | None | None | >90 | 9.8 | |
| **H6** | 47 | M | B | 81.9<br>180<br>25.3 | Healthy | 13 pack yrs; active | None | >90 | 8.1 | |
| **H7** | 62 | M | W | 74<br>178<br>23.4 | Healthy | None | None | >90 | 9.1 | *IPF control |
| **Pulmonary fibrosis subjects** | | | | | | | | | | |
| **IPF1** | 70 | M | W | 71.4<br>164<br>27 | IPF/UIP | 4 pack yrs, remote | Nintedanib | 102 | 8.3 | |
| **IPF2** | 57 | F | W | 55.2<br>154<br>23 | Familial IPF/ UIP | None | Pirfenidone | 59 | 9.3 | |
| **IPF3** | 75 | M | W | 74.4<br>176<br>23.5 | IPF/RA | 35 pack yrs, none for 18 yrs | Pirfenidone Abatacept | 92 | 8.1 | |
| **IPF4** | 62 | M | W | 100<br>183<br>29.9 | IPF/UIP | 50 pack yrs, none for 6 years | Pirfenidone | 79 | 7.1 | |

**Abbreviations:** F, female; M, male; B, black; W, White; Wt; weight; Ht, height; BMI, Body mass index; IPF, Idiopathic pulmonary fibrosis; UIP, usual interstitial pneumonia; PPD, packs of cigarettes per day; FVC, Forced vital capacity as percent predicted; PF, Pulmonary fibrosis; RA, Rheumatoid arthritis.
*Dynamic PET imaging as a control for pulmonary fibrosis, not included in dosimetry study.

**Table S4. Estimated human dosimetry**

| Organ | Female dose (rad/mCi) | Male dose (rad/mCi) |
|---|---|---|
| Adrenals | 0.044 | 0.027 |
| Brain | 0.027 | 0.016 |
| Breasts | 0.027 | - |
| Gallbladder wall | 0.049 | 0.031 |
| Lower large intestine wall | 0.094 | 0.048 |
| Small intestine wall | 0.037 | 0.024 |
| Stomach wall | 0.035 | 0.022 |
| Upper large intestine wall | 0.065 | 0.034 |
| Heart wall | 0.079 | 0.047 |
| Kidneys | 0.412 | 0.306 |
| Liver | 0.242 | 0.132 |
| Lungs | 0.033 | 0.019 |
| Muscle | 0.026 | 0.021 |
| Ovaries | 0.040 | - |
| Pancreas | 0.042 | 0.026 |
| Red marrow | 0.028 | 0.018 |
| Osteogenic cells | 0.060 | 0.034 |
| Skin | 0.025 | 0.015 |
| Spleen | 0.056 | 0.034 |
| Testes | - | 0.021 |
| Thymus | 0.030 | 0.018 |
| Thyroid | 0.027 | 0.017 |
| Urinary bladder wall | 0.567 | 0.471 |
| Uterus | 0.045 | - |
| Total Body exposure | 0.043 | 0.028 |
| Effective Dose Equivalent (rem/mCi) | 0.107 | 0.075 |
| Effective Dose (rem/mCi) | 0.077 | 0.053 |
